# Clustering Sperm: A Statistical Approach to Identify Sperm Morph Numbers in the *Drosophila Obscura* Species Group

**DOI:** 10.1101/2025.05.29.656784

**Authors:** Fiona Messer, Helen White-Cooper

## Abstract

The *Drosophila obscura* species group is sperm heteromorphic. Sperm heteromorphism is production of multiple sperm types, or ‘morphs’, within a single male. In the case of the *obscura* species, males produce two or three sperm morphs of different sizes, simultaneously, within the same testis, and throughout their lifetime. A long sperm morph, the ‘eusperm’ is fertile, whereas shorter sperm morphs, the ‘parasperm’, are non-fertilising and protect the eusperm within the female reproductive tract. Many studies over the past 55 years have measured sperm length of eusperm and parasperm in *obscura* species. One recent study found that *D. pseudoobscura* produces three sperm morphs, two parasperm and one eusperm, in contrast to the two morphs described in older studies of this species. Thus, the number of morphs made by some species in this clade is still in dispute. Here, we review sperm length data from previous studies and re-measure sperm of eight species including *D. pseudoobscura*. We used two statistical cluster analysis approaches to identify sperm morphs based on sperm and nucleus length, to test whether multiple parasperm morphs are present in more species than previously identified. We confirmed the presence of two parasperm morphs in *D. pseudoobscura*, and found that two closely related species, *D. persimilis* and *D. miranda*, also produce two parasperm morphs. *D. affinis, D. azteca, D. bifasciata, D. guanche* and *D. subobscura* produce a single parasperm morph, indicating that the presence of two parasperm is a derived feature in the *pseudoobscura* species subgroup.

## 1. Introduction

### 1.1. Sperm heteromorphism in the *obscura* species group

Sperm heteromorphism is the production of multiple sperm morphs by an individual male. Morphs may be produced simultaneously or sequentially, and within the same testis or in separate testes. Morphs are produced consistently between males and are not the result of developmental aberration. Sperm morphs are distinct in morphology and function. For example, they may vary in size and/or DNA content. One or more morphs is non-fertilising, having some other function in fertility and reproduction (Bernasconi and Hellriegel, 2005, Snook et al., 1994, Snook and Karr, 1998, Swallow and Wilkinson, 2002).

The *obscura* species group of *Drosophila* produce multiple morphs of differing sizes, a form of sperm heteromorphism sometimes termed ‘polymegaly’ (Beatty and Sidhu, 1970). In these species, a longer morph – the ‘eusperm’ – is fertile, while one or more shorter morphs – the ‘parasperm’ – are non-fertilising. Shorter parasperm protect the longer eusperm from female-mediated spermicide present in the female reproductive tract (Holman and Snook, 2008, Alpern et al., 2019). The number of morph classes and their functions have been debated since the phenomenon was first described in the *obscura* species by Beatty and Sidhu (1970). This is discussed in detail below, by species subgroup. Table 1 summarises published sperm morph lengths by species.

**Table 1.**
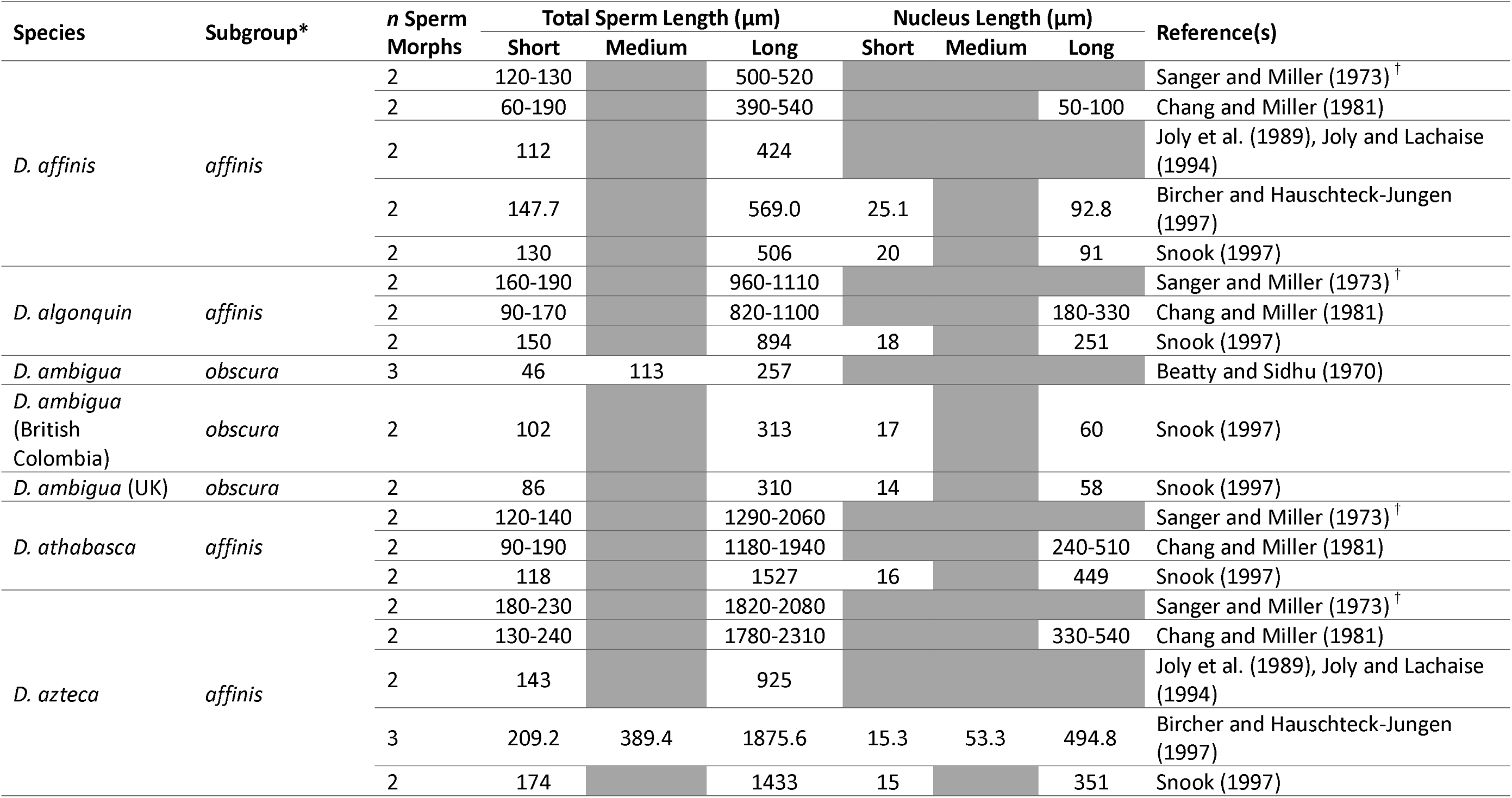
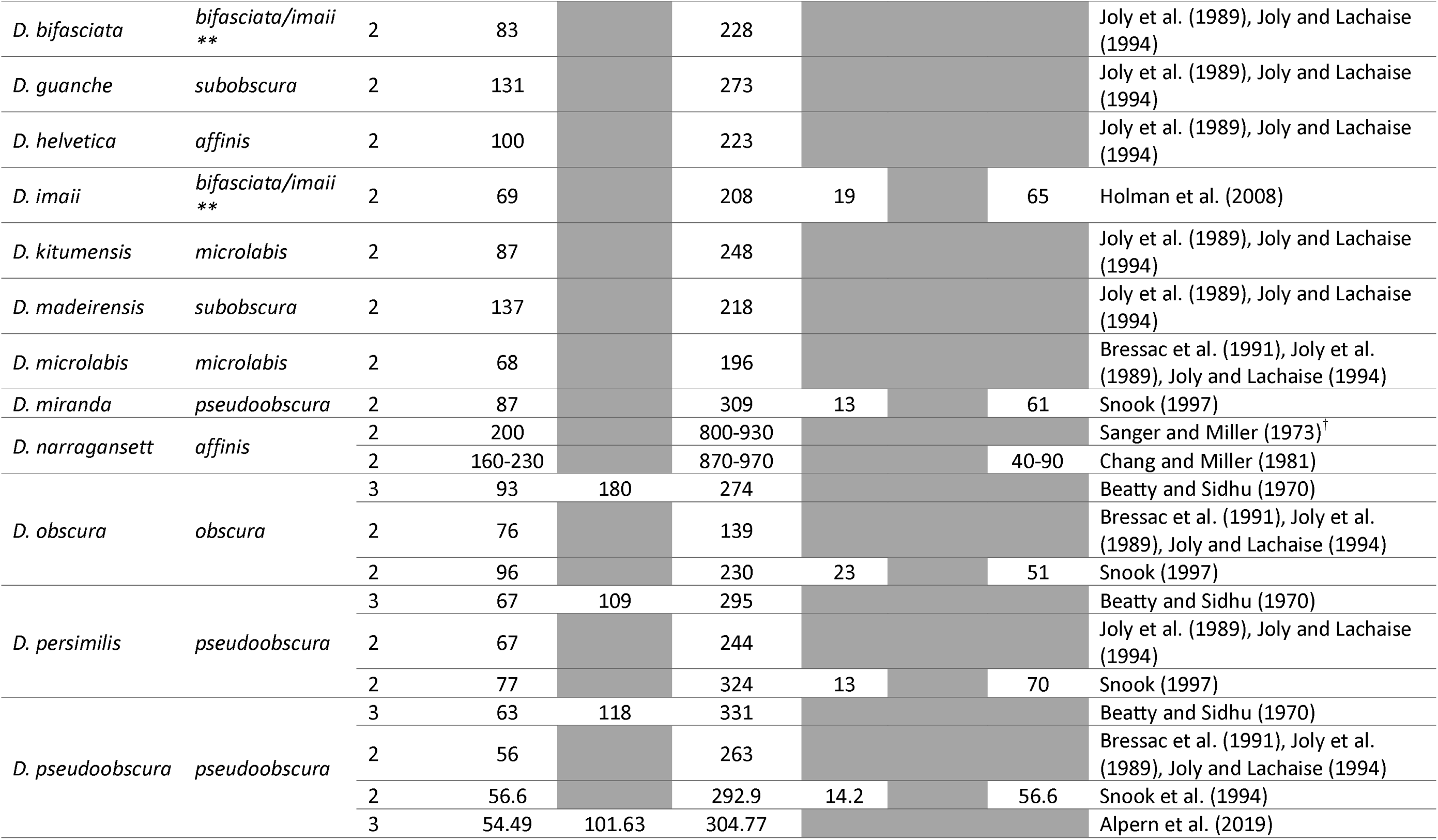
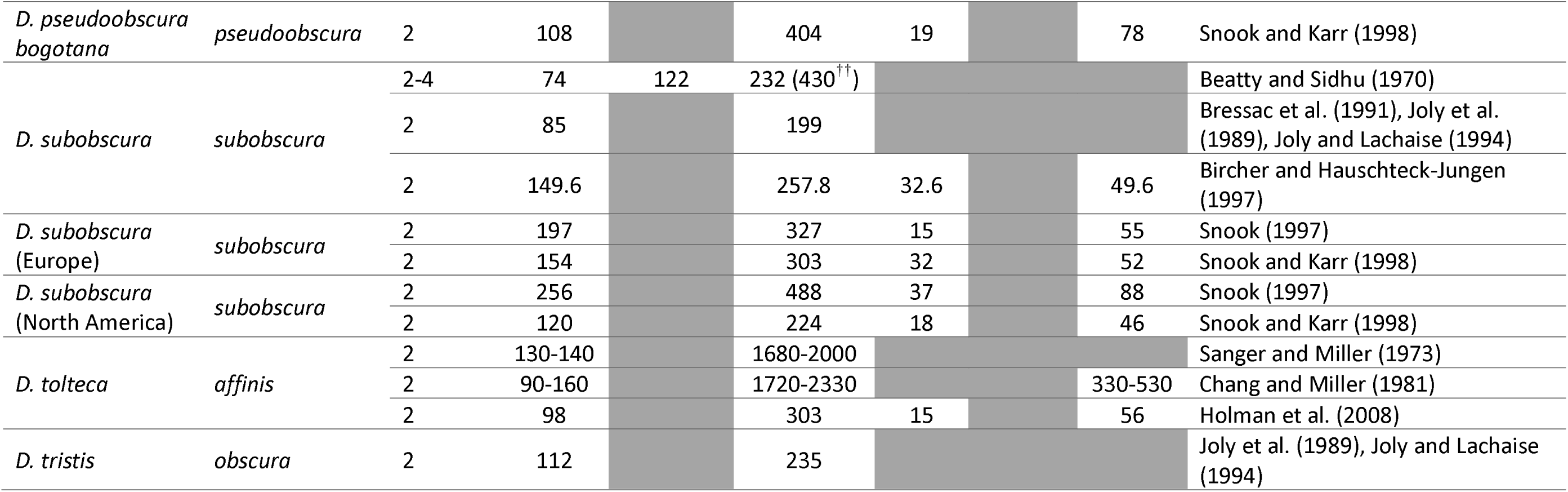
– Summary of the number of sperm morphs identified, and total lengths and nucleus lengths of heteromorphic sperm in the obscura species group in published literature. *Species subgroups according to O’Grady (1999), Barrio and Ayala (1997), Gao et al. (2007). **Older literature groups D. bifasciata and D. imaii with the obscura species, however some more recent studies have found the bifasciata/imaii and obscura subgroups to be paraphyletic. *^†^*Sanger and Miller (1973) measured multiple populations for each species, however data is shown as a range as all populations were from similar geographical distributions. *^††^*430µm sperm in D. subobscura observed once.

#### 1.1.1. affinis – D. affinis, D. azteca, D. algonquin, D. athabasca, D. helvetica, D. tolteca and D. narraganset

The *affinis* subgroup was first investigated by Sanger and Miller (1973) then later by Joly et al. (1989), finding two classes in all seven species. The *affinis* subgroup has the greatest variability between species ranging from parasperm lengths of 100μm in *D. helvetica* to 240μm in *D. azteca*, and eusperm lengths of 220μm up to 2310μm, again in *D. helvetica* and *D. azteca*, respectively (Joly et al., 1989, Chang and Miller, 1981).

A later study by Bircher and Hauschteck-Jungen (1997) utilised cluster analysis (method not specified) of sperm length to identify the number of morph classes in both *D. affinis* and *D. azteca*. They found two morphs in *D. affinis*, but three in *D. azteca*, although they note that the medium morph in *D. azteca* was observed in only two of the five males.

#### 1.1.2. bifasciata – D. bifasciata and D. imaii

*D. bifasciata* and *D. imaii* sperm morphs have each been measured once. Both species produce two morphs, 83 and 228μm in *D. bifasciata* and 69 and 208μm in *D. imaii*. (Joly et al., 1989, Joly and Lachaise, 1994, Holman et al., 2008)

#### 1.1.3. obscura – D. obscura, D. ambigua and D. tristis

*D. obscura* and *D. ambigua* were both included in Beatty and Sidhu’s 1970 study (Beatty and Sidhu, 1970). Three morph classes were identified in *D. obscura*, 93, 180 and 274µm in length and *D. ambigua*, 46, 113, and 257µm in length. Subsequent studies have not identified a ‘medium’ class, finding only two morph classes in both species. *D. tristis* were also found to have two morph classes, 112 and 235µm in length (Joly et al., 1989).

*D. obscura* parasperm has been measured to be between 76-96µm, consistent with earlier studies (Joly et al., 1989, Bressac et al., 1991, Joly and Lachaise, 1994). However, eusperm length varies considerably between these studies, 139µm in Joly et al. (1989), and 230µm in Snook (1997), closer to that found by Beatty and Sidhu (1970). This variation in eusperm length may be due to naturally occurring population variation, as has been observed in other species (Snook, 1997, Snook and Karr, 1998, Joly and Lachaise, 1994).

Snook (1997) measured sperm and nucleus length in two populations of *D. ambigua*, noting a small difference in lengths between UK and Canadian populations. In both cases, two morphs were identified, 86-102µm and 310-313µm.

#### 1.1.4. pseudoobscura – D. pseudoobscura, D. persimilis *AND* D. miranda

Arguably the most studied species within the *obscura* species group, *D. pseudoobscura* were originally described as having three sperm morphs: a short 63µm morph, a medium 118µm morph, and a long 331µm morph (Beatty and Sidhu, 1970, Beatty and Burgoyne, 1971). *D. persimilis* was also described as having three morphs, 67µm, 109µm and 295µm in length, although the shorter morphs were considered ‘indistinctly separate’ (Beatty and Sidhu, 1970, Beatty and Burgoyne, 1971). *D. pseudoobscura* and *D. persimilis* were revisited in a larger study by Joly et al. (1989) and Joly and Lachaise (1994), which found a bimodal distribution of morphs in both species. A small number of ‘extra-long’ sperm were observed in *D. pseudoobscura*, although this was not further discussed (Joly and Lachaise, 1994). Snook et al. (1994) found evidence for only two sperm morphs in *D. pseudoobscura*, although did note that there was a possibility for the presence of two ‘short’ parasperm size classes, which would be difficult to distinguish. Sperm head (nuclear) length was also characterised, finding distinct head length for the two size classes identified (parasperm = 14.2µm, eusperm = 56.6µm).

*D. pseudoobscura* was revisited again by Alpern et al. (2019), who found further evidence in support of earlier observations by Beatty and Sidhu (1970) that there are three sperm morphs produced in this species. They identified two distinct parasperm morphs, for which the lengths were non-overlapping; 55μm parasperm 1 and 102μm parasperm 2. They used a statistical approach to confirm that the three groups were significantly different from each other, based on total sperm length (ANOVA with Tukey’s post-hoc test). Morphological differences between the parasperm morphs were observed: a ‘string-like structure’ associated with the tails of parasperm 2 and eusperm, which appears to be required for the spiral conformation of the tail of these two morphs but was not observed in parasperm 1.

*D. miranda* was not discussed in the earlier studies of *Drosophila* sperm heteromorphism, but was later described as having two morphs, 87μm and 309μm (Snook, 1997).

#### 1.1.5. microlabis – D. microlabis and D. kitumensis

*D. microlabis* and *D. kitumensis* have each been measured once. Both produce two morph classes. *D microlabis* has 68µm parasperm and 196µm eusperm. *D kitumensis* has 87µm parasperm and 248µm eusperm.

#### 1.1.6. subobscura – D. subobscura, D. guanche *AND* D. madeirensis

*D. subobscura* was initially suggested to have up to four morphs by Beatty and Sidhu (1970), finding 74µm short, 122µm medium, 232µm long, and 430µm extra-long sperm, although only one 430µm was observed in that study. *D. subobscura* has been re-measured more than any other species, across multiple populations and geographic distributions, and a substantial variation in both parasperm and eusperm lengths is apparent. Parasperm length varies between 85-256µm, with eusperm length between 199-488µm (Joly et al., 1989, Bressac et al., 1991, Joly and Lachaise, 1994, Bircher and Hauschteck-Jungen, 1997, Snook, 1997, Snook and Karr, 1998, Beatty and Burgoyne, 1971). In cluster analysis of *D. subobscura* sperm length, Bircher and Hauschteck-Jungen (1997) found two morph classes, as have all other studies subsequent to Beatty and Sidhu (1970).

*D. guanche* and *D. madeirensis* are less-well studied than *D. subobscura*. Both have two morph classes (Joly et al., 1989, Joly and Lachaise, 1994).

### 1.2. Inter– and intra-specific variation in sperm length

Joly and Lachaise (1994) analysed sperm length variance within and between males, and between strains of *D. affinis*, finding significant differences in the length of long sperm between individuals and populations, and in the length of short sperm between individuals within populations. They also compared sperm length between closely related species (within species subgroups), finding greater interspecific variation in eusperm length compared to parasperm, based on coefficient of variation.

Broadly, *Drosophila* sperm length is highly correlated with the length of the female sperm storage organs, the spermathecae (Pitnick et al., 1999). In *obscura* species specifically neither eusperm length nor parasperm length were correlated to spermathecal area. However, eusperm length, but not parasperm length, is correlated with the length of the female ventral receptacle, suggesting the morphs have evolved differing functions (Holman et al., 2008).

### 1.3. Relative proportions of sperm morphs

The proportion of eusperm to parasperm within the ejaculate can be measured by several different methods:

1. Individualising spermatid cysts
2. Sperm extracted from seminal vesicle (‘produced’)
3. Sperm extracted from female reproductive tract (‘transferred’)

Published data on parasperm proportion in ejaculate can be found in Table 2. Despite different methodologies and sample populations, proportion remains relatively consistent between studies. Proportion does change as the male ages; one-day-old *D. subobscura* males have almost no eusperm in the seminal vesicle, whereas seminal vesicles of four-day old males contain both morphs (Bircher et al., 1995). This suggests that eusperm require more time to develop than parasperm. There may also be an effect of rearing stocks in the lab. In comparison to wild-caught populations, laboratory-reared *D. pseudoobscura* transfer an increased proportion of eusperm (Snook and Markow, 2002).

**Table 2:**
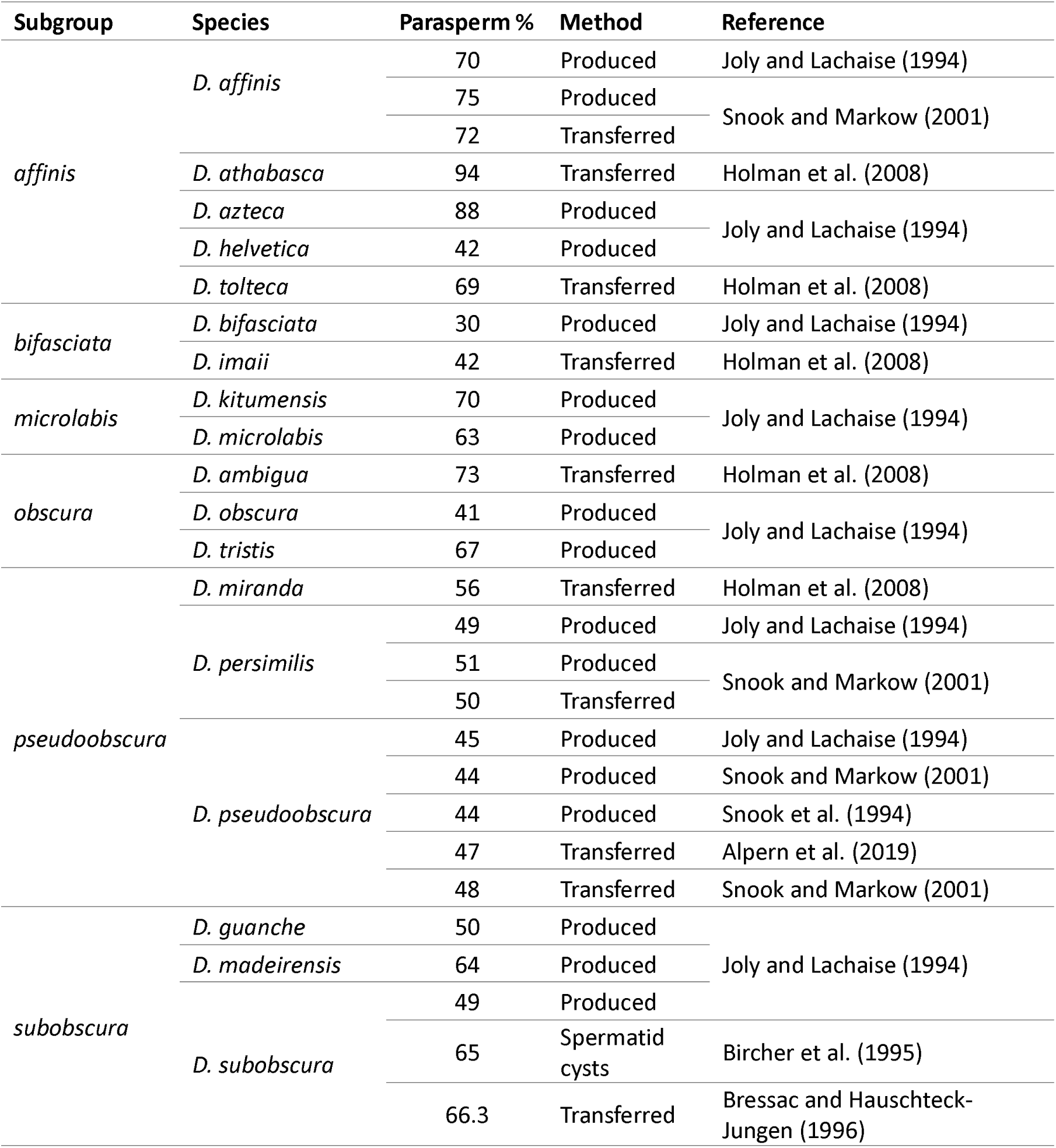
Proportion of parasperm in the ejaculate of obscura group species, by species subgroup. Method of sperm extraction is specified. Produced = sperm dissected from seminal vesicle. Transferred = sperm dissected from female reproductive tract. Spermatid cyst = individualising sperm cysts dissected from testis.

Between species, the proportion of parasperm in the ejaculate correlates with eusperm length. Parasperm comprise a greater proportion of the ejaculate in species with longer eusperm (Holman et al., 2008).

### 1.4. Functions of parasperm in the *obscura* species group

Both eusperm and parasperm contain an equal amount of DNA, however only eusperm, and never parasperm, are observed in the fertilised eggs of *D. pseudoobscura* suggesting that only eusperm are capable of fertilisation (Snook et al., 1994). This analysis was later expanded to over 1500 eggs from *D. affinis*, *D. athabasca, D. miranda*, *D. persimilis* and *D. subobscura*, which were also found to contain only eusperm sperm (Snook and Karr, 1998). Snook and Karr (1998) suggested that there may be a barrier to parasperm entry into egg via sperm-egg interactions, such as a lack of gamete surface membrane receptors, or parasperm heads being too wide to enter the micropyle.

Instead of fertilisation, parasperm have a protective role. The female reproductive tract is a spermicidal environment, in which eusperm viability declines over time. When incubated with female reproductive tract tissue extract, increasing the proportion of parasperm in the ejaculate increases eusperm survival. Males which transfer a higher proportion of parasperm in the ejaculate also show higher eusperm viability after 30 minutes in the female reproductive tract. This indicates that parasperm protect eusperm in the female reproductive tract (Holman and Snook, 2008, Alpern et al., 2019).

Where multiple parasperm morphs are present, the medium parasperm 2 may also have a function in sperm competition. Alpern et al. (2019) found increased sperm competition increases the percentage of parasperm 2 in the ejaculate, while eusperm and parasperm 1 percentage decreases. They suggest that parasperm 2 may remove sperm of competing males already present in the female reproductive tract and spermathecae.

### 1.5. Statistical methods for analysis of sperm morph classes

While most studies have used a visual approach to assigning morph classes, statistical methods to determine the number of sperm morphs, and their significance, have been used in some previous studies of sperm heteromorphism. Snook (1995) used an approach of testing sperm length distributions for normality, then describing the shape of the distribution by kurtosis. Kurtosis indicated most species in the *obscura* species group produce two morphs, but for *D. pseudoobscura*, *D. miranda* and a European population of *D. ambigua*, kurtosis indicated two parasperm morphs – short and medium, although these were not discrete. Alpern et al. (2019) used ANOVA to test for significance in difference of sperm lengths between putative morphs in *D. pseudoobscura*. Short, medium and long morph total sperm length was significant by Tukey’s post-hoc. They also found a significant difference in nucleus length between parasperm 1 and 2.

In this study we use a cluster analysis approach to group sperm into morph classes based on total length and nucleus length. We used two mathematically dissimilar clustering methods, hierarchical cluster analysis and Gaussian mixture modelling, and compare the results of both models.

### 1.6. Aim of study

The aim of this study is to reanalyse sperm length and the number of sperm morph classes in eight species of the *obscura* species group: *D. affinis*, *D. azteca*, *D. bifasciata*, *D. guanche*, *D. miranda*, *D. persimilis*, *D. pseudoobscura* and *D. subobscura*. All eight have previously been analysed, however recent evidence of multiple parasperm morphs in *D. pseudoobscura* gives reason to reassess the number of morphs present in other species, particularly those to which it is most closely related – *D. persimilis* and *D. miranda*. We use a new statistical approach to identify morphs, based on cluster analysis of sperm morphological data. We also discuss whether there is evidence of multiple eusperm morphs in *obscura* group species.

## 2. Methods

### 2.1. Fly Stocks

*D. pseudoobscura* SLoB3 stocks were gifted by T. Price (University of Liverpool, UK). *D. miranda* stocks were gifted by D. Bachtrog (UC Berkeley, USA). All other stocks were obtained from the National Drosophila Species Stock Center (Cornell University, USA) and Kyorin-Fly Stock Center (Kyorin University, Japan) (Table 3).

**Table 3.**
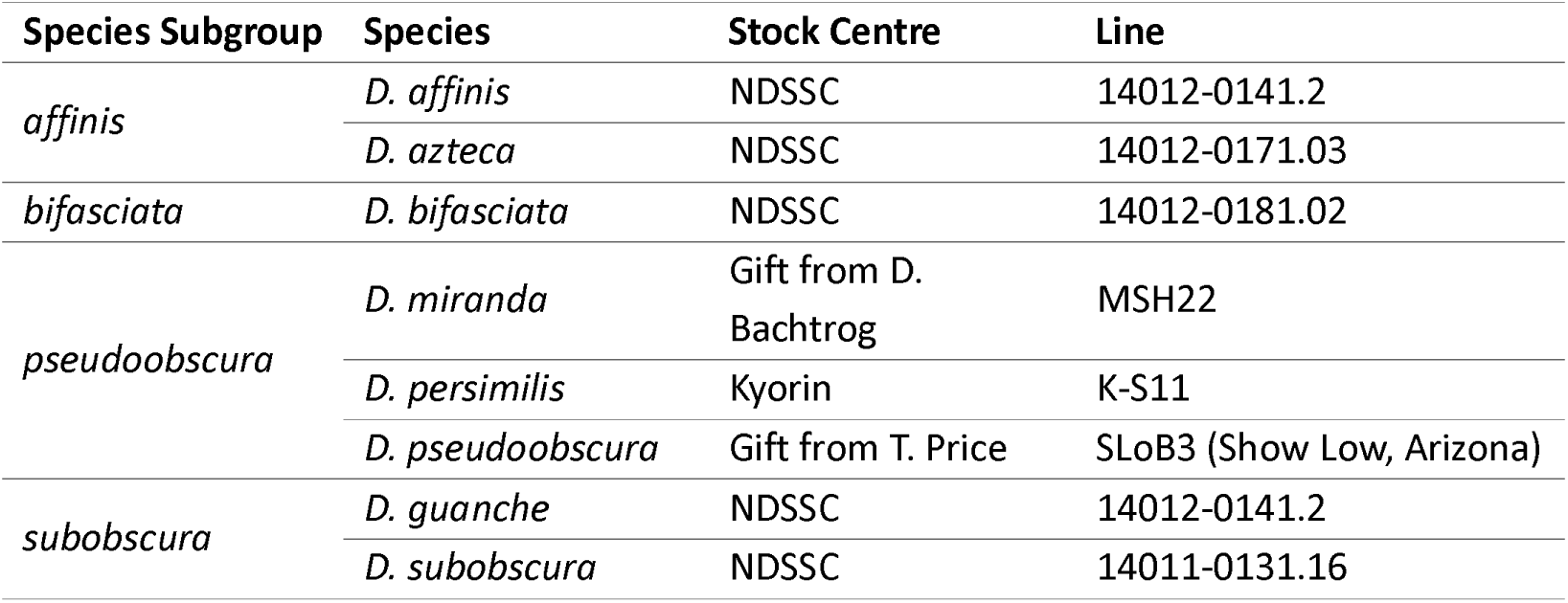
obscura species group stocks used for sperm isolation, imaging, and measurements.

| Species Subgroup | Species | Stock Centre | Line |
| --- | --- | --- | --- |
| *affinis* | *D. affinis* | NDSSC | 14012-0141.2 |
|  | *D. azteca* | NDSSC | 14012-0171.03 |
| *bifasciata* | *D. bifasciata* | NDSSC | 14012-0181.02 |
| *pseudoobscura* | *D. miranda* | Gift from D. Bachtrog | MSH22 |
|  | *D. persimilis* | Kyorin | K-S11 |
|  | *D. pseudoobscura* | Gift from T. Price | SLoB3 (Show Low, Arizona) |
| *subobscura* | *D. guanche* | NDSSC | 14012-0141.2 |
|  | *D. subobscura* | NDSSC | 14011-0131.16 |

### 2.2. Sperm dissection and fixation

Testes were dissected into drop of testis buffer (183mM KCl, 47mM NaCL, 10mM Tris HCl, pH6.8) with seminal vesicles attached. Seminal vesicles were separated from the testes then transferred to a fresh drop of testis buffer. To remove sperm from the seminal vesicles, the seminal vesicles were held with a pair of forceps at one end, and the sperm squeezed out by running a fine tungsten needle along the length of the seminal vesicle. Empty seminal vesicles were discarded. Sperm was separated in testis buffer by gently mixing with a fine tungsten needle. 20µL of sperm in testis buffer was transferred to a 1.5mL tube. The mixture was gently pipetted 5X then incubated at room temperature for 5 minutes, to allow further dissociation. To fix, 20µL 4% paraformaldhyde in PBS+0.1% Tween 20 was added to the sperm/testis buffer mixture and incubated at room temperature for 5 minutes. 40µL mounting medium (80% glycerol, 2.5% n-propyl gallate, 1µg/mL Hoechst 33258) was added to the fixed sperm and mixed by pipetting 5X. 30µL of the fixed sperm/mounting medium mixture was transferred to a microscope slide and a coverslip added.

A minimum of four males and 200 sperm were counted from each species (see S1 for data).

### 2.3. Imaging and Image Analysis

Slides were imaged using the Olympus BX50 (Olympus Europa, Germany) with Hamamatsu ORCA-05G camera attachment and HCImage software (v.2,2,6,4 Hamamatsu Corp. 2011). Each sperm was imaged by brightfield and fluorescence (405nm) for whole-sperm and nucleus, respectively. Imaging was performed in transects across the slide to ensure unbiased sampling of the sperm population. Measurements were performed with the ImageJ Measurement tool (ImageJ 1.54f, NIH). Nucleus measurements were taken from the base of the nucleus to the tip of the sperm head.

### 2.4. Statistical Analysis

All statistical analysis was performed with R Studio v.4.2.1 (R Core Team, 2022) (see S2 for R scripts, S3 for model outputs). Cluster analysis was performed using the total sperm length and nucleus length measurements, both measured in microns. The data was not scaled prior to analysis.

Hierarchical cluster analysis (HCA) was performed with the ‘hclust’ function of the ‘stats’ package (R Core Team, 2022). The optimum number of clusters was first determined by gap statistic modelling (method = first maximum, bootstrap = 1000) (Tibshirani et al., 2001) using the ‘cluster’ package (Maechler et al., 2022). Total sperm length and nucleus length data was transformed into a distance matrix using the Euclidean distance. Hierarchical clustering was performed using the Ward distance. Clustering was assigned using the ‘cutree’ function, with the number of clusters specified by the gap statistic. The distribution of total sperm length within clusters assigned by HCA was tested for normality by Shapiro-Wilk (S4).

Gaussian mixture modelling (GMM) was performed with the ‘mclust’ package (Scrucca et al., 2016). GMM estimates the number of clusters and assigns datapoints to clusters by model fitting, so no other methods were required to estimate the number of clusters prior to running the model.

All scatterplots and dendrograms were drawn with ‘ggplot2’ (Wickham, 2016) and ‘ggdendro’ (de Vries and Ripley, 2022) packages.

The proportion of parasperm within the ejaculate was calculated as:

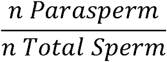

The proportion of parasperm length within the total sperm length produced was calculated as:

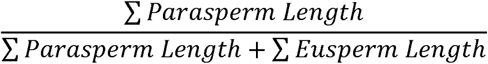

## 3. Results

### 3.1. Summary of results

Data for each species was plotted on a scatterplot of total sperm length against nucleus length (*Error! Reference source not found.*). As has previously been observed, more than one sperm morph was present in all eight species studied. There was substantial variation between species in total sperm and nucleus lengths. The two cluster analysis methods produced different results in several species (Table 4). Gaussian mixture modelling often identified a greater number of clusters than HCA, suggesting it is a more sensitive method and/or is more prone to false positives. The number of sperm morphs, mean total sperm lengths, nucleus lengths, standard deviations, and proportion each morph contributes to the total sperm produced for each species are summarised in Table 5. The proportion of total sperm length consisting of parasperm was variable between species, between 0.08-0.42, with a mean of 0.25 (S5)

**Table 4:** Summary of number of clusters identified in sperm morphological data from obscura group species. Two analysis methods were used, hierarchical cluster analysis and Gaussian mixture modelling.

| Species Subgroup | Species | *n* Clusters |  |
| --- | --- | --- | --- |
|  |  | Hierarchical Cluster Analysis | Gaussian Mixture Modelling |
| *affinis* | *D. affinis* | 2 | 3 |
|  | *D. azteca* | 3 | 3 |
| *bifasciata* | *D. bifasciata* | 3 | 3 |
| *pseudoobscura* | *D. miranda* | 3 | 4 |
|  | *D. persimilis* | 2 | 5 |
|  | *D. pseudoobscura* | 3 | 6 |
| *subobscura* | *D. guanche* | 2 | 4 |
|  | *D. subobscura* | 2 | 2 |

**Table 5:**
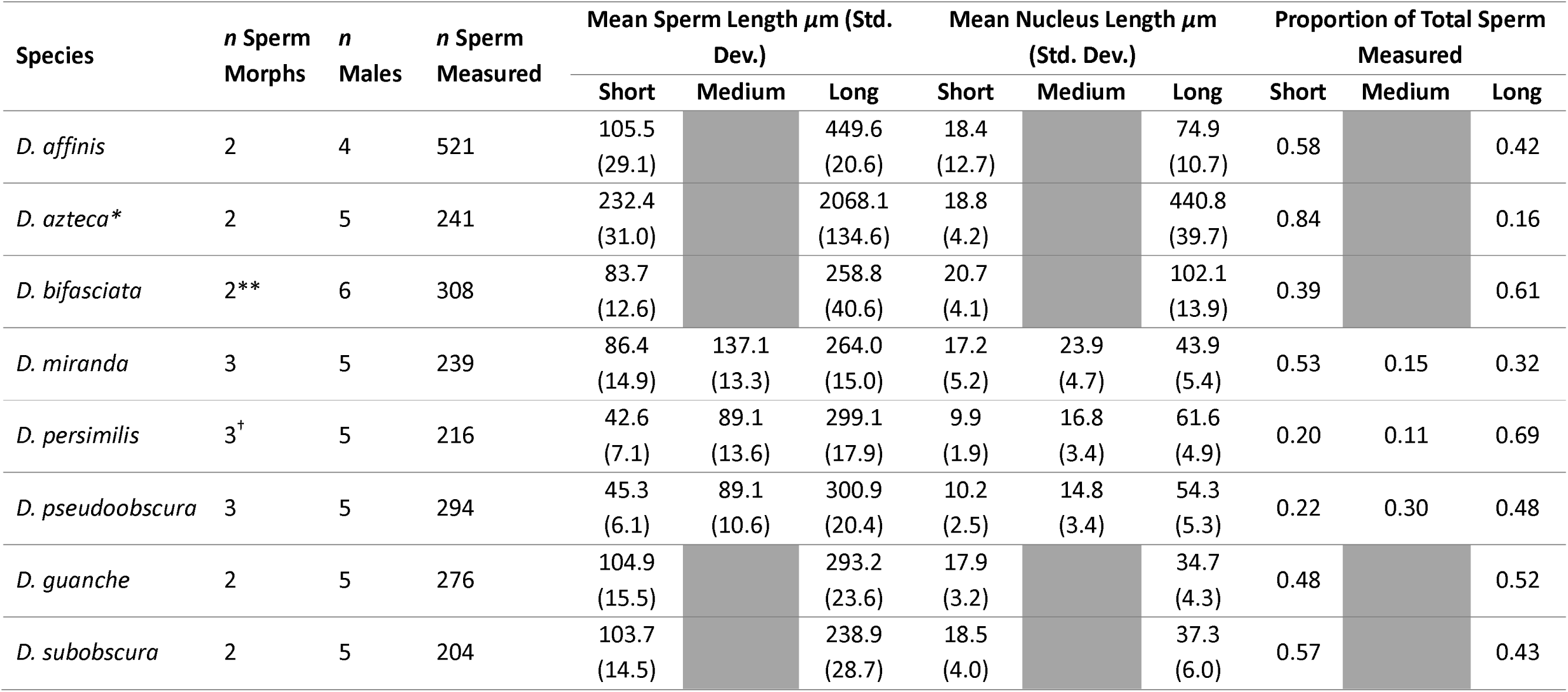
Means of sperm morphs based on hclust clustering. *D. azteca broken eusperm cluster was removed prior to calculating total sperm and nucleus length means, proportion of total sperm measured included broken eusperm in the ‘long’ cluster. Note that eusperm were not randomly sampled from D. azteca, therefore calculations for proportion of total sperm measured are not representative of proportion within the ejaculate. **D.bifasciata hclust and GMM clustering identified two eusperm clusters, however clusters appeared to be resulting from individual variation. Two eusperm clusters were combined for mean total sperm length, nucleus length and proportion. *^†^*D. persimilis hclust identified two clusters, GMM identified an additional parasperm cluster which is supported by morphological evidence.

In addition to length, there are other morphological differences between eusperm and parasperm morphs (Figure 2). Eusperm nuclei are long, and taper to a thin point at the tip of the sperm, whereas parasperm nuclei appear more rounded at the tip, and taper towards the tail of the sperm. When imaged together, there is often a clear difference in the brightness of the DNA stain between the morphs, with parasperm nuclei being shorter and more compact, and thus appearing brighter. Parasperm nuclei often appeared hook-or ‘C’-shaped compared to more linear eusperm nuclei, and parasperm tails were often more tightly coiled in appearance, while eusperm tails were wavier.

**Figure 1:**
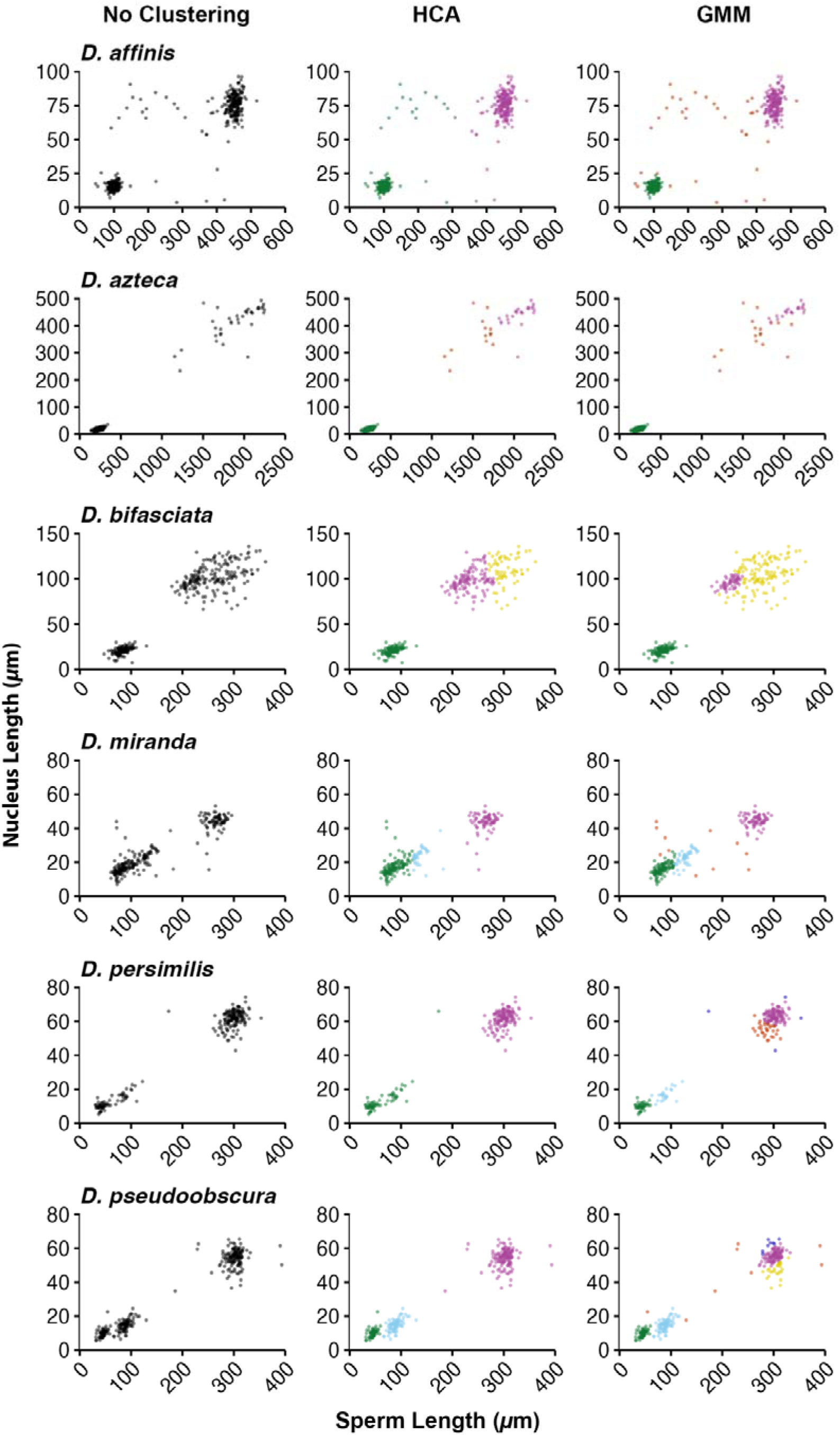
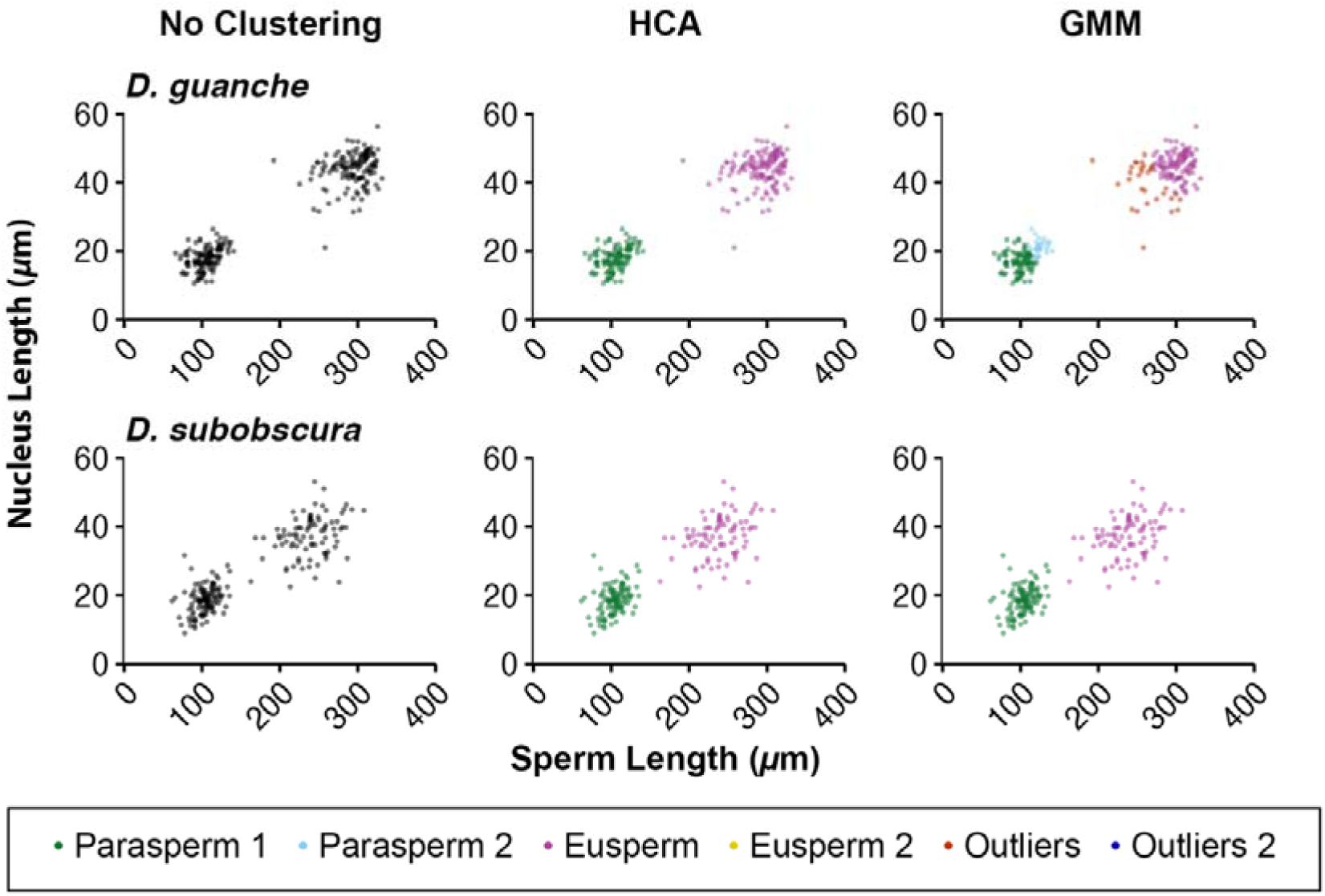
Scatterplots of total sperm length against nucleus length. For each species, data is shown with no clustering, clustering according to hierarchical cluster analysis (hclust) and Gaussian mixture modelling (mclust). Note different X and Y axis scales between species.

**Figure 2:**
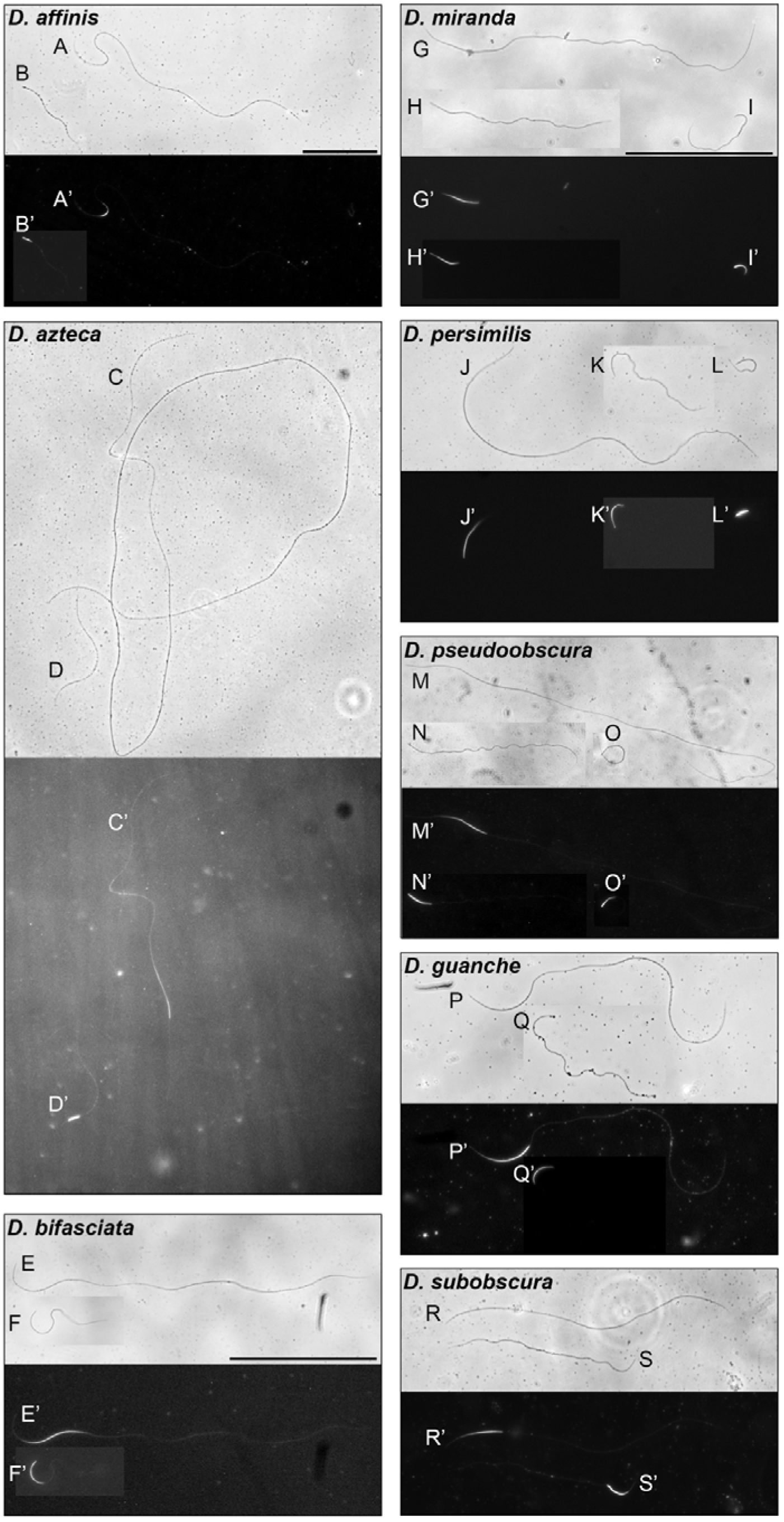
Mature spermatozoa from Drosophila obscura group species. A-S: Brightfield imaging of whole sperm. A’-S’: Fluorescence imaging of whole sperm showing nuclei stained with Hoechst 33258. A, C, E, G, J, M, P and R: Eusperm. B, D, F, Q and S: Parasperm. H, K and N: Parasperm 2, or medium sperm, from D. miranda, D. persimilis, and D. pseudoobscura. I, L and O: Parasperm 1, from D. miranda, D. persimilis and D. pseudoobscura. Note the long needle-shaped nuclei of eusperm compared to shorter parasperm nuclei, often hook or ‘C’ shaped. Brightfield images brightness has been increased. D. azteca fluorescence image brightness and contrast increased. Scale = 100µm.

### 3.2. affinis – D. affinis and D. azteca

#### 3.2.1. D. affinis

*D. affinis* sperm was clustered into two clusters by HCA, with a further cluster identified by GMM of outliers likely to be mostly broken eusperm. By HCA, neither parasperm nor eusperm lengths were normally distributed with outliers (parasperm W = 0.500, p < 0.001; eusperm W = 0.859, p < 0.001), but were normally distributed when the outliers were removed from the dataset (parasperm W = 0.993, p = 0.251; eusperm W = 0.993, p = 0.372). Mean parasperm length was 105.5µm, mean eusperm length was 449.5µm, and parasperm comprised 58% of the sperm total sperm measured (Figure 2A-B).

#### 3.2.2. D. azteca

Both HCA and GMM identified three clusters in *D. azteca*, one parasperm cluster, and two eusperm clusters. However, like *D. affinis*, it is likely that broken eusperm make up the shorter of the two eusperm clusters, as this cluster contains fewer individual sperm and shows large variation in total sperm length – between 1155 and 1743µm. *D. azteca* has the longest eusperm of the *obscura* species group and therefore is more likely to break during dissection. Mean parasperm length was 232.4µm. Mean eusperm length when broken sperm were removed from the dataset was 2068.1µm (Figure 2C-D). Both parasperm and eusperm length, with broken eusperm removed from the data, were normally distributed (parasperm W = 0.992, p = 0.301; eusperm W = 0.934, p = 0.106).

Of the total sperm measured 84% were parasperm, and 16% eusperm. However, due to the small numbers of eusperm produced by *D. azteca*, we measured all eusperm from each individual rather than a random transect of the slide as was done for all other species. Therefore, eusperm proportion of the total sperm measured is not representative of the eusperm proportion within the ejaculate for *D. azteca*.

### 3.3. D. bifasciata

Both HCA and GMM identified three clusters in D. bifasciata, one parasperm cluster and two eusperm clusters. Mean parasperm length was 83.7µm and parasperm comprised 39% of the total sperm measured. Parasperm length was normally distributed (W = 0.988, p = 0.356).

Whereas in *D. affinis* and *D. azteca*, the third cluster was more obviously broken eusperm, this does not appear to be the case in *D. bifasciata*. *D. bifasciata* eusperm are much shorter than those of the *affinis* species subgroup, at a mean length of 258.8µm, therefore less likely to break during preparation. When the two HCA eusperm clusters were combined the data were not normally distributed (W = 0.97, p = 0.000932), indicating a bimodal distribution. However, when separated neither the eusperm 1 (W = 0.97, p = 0.021) nor eusperm 2 (W = 0.95, p = 0.014) HCA clusters were normally distributed, indicating that the clusters are not distinct. The question therefore is – does *D. bifasciata* produce two eusperm morphs, or is eusperm simply more variable in length than is observed in other species?

When clustered by HCA, the two eusperm clusters were present in all the individuals measured (although this was not the case for GMM clustering). However, while both models identify two eusperm clusters, the cluster identity of individual datapoints is not consistent between the two models – 35% of eusperm morphs were placed in a different cluster in the GMM analysis compared to HCA. The mean lengths of the eusperm clusters, by HCA, was 234.4 and 304.4µm, and by GMM was 215.4 and 277.7µm. Mean nucleus length by HCA was 98.1 and 109.6µm, and by GMM was 94.9 and 103.9µm. Nucleus length is notable for its similarity between the two clusters, in both models. Furthermore, other than length, there were no clear morphological differences between eusperm clusters. There were considerable differences between individuals in the range of sperm lengths observed (S6). Taken together, this indicates that the two eusperm clusters are not distinct and it is not possible to determine whether these are distinct morphs. Further morphological and functional analysis would be required to confirm this.

### 3.4. pseudoobscura – D. miranda, D. persimilis and D. pseudoobscura

#### 3.4.1. D. miranda

HCA identified three clusters in *D. miranda;* two parasperm clusters and one eusperm cluster. GMM identified an additional cluster which is likely to be mostly broken eusperm. Mean eusperm length was 264µm. Eusperm length was normally distributed (W = 0.980, p = 0.280).

Mean parasperm 1 length was 89.4µm whereas mean parasperm 2 length was 137.1µm. Neither parasperm 1 nor parasperm 2 length were normally distributed (parasperm 1 W = 0.950, p < 0.001; parasperm 2 W = 0.804, p < 0.001). Parasperm length was also not normally distributed when the clusters were combined (W = 0.906, p < 0.001), indicating that the clusters are separate, but not as distinct as *D. pseudoobscura* (see below). The two parasperm clusters as identified by GMM were present in all individuals from which sperm were measured (by HCA clustering, none of the 27 sperm measured from male 2 were parasperm 2).

Morphological differences were observed between the two parasperm clusters. While the tails of both morphs were similarly coiled, the head and nucleus of parasperm 1 was often hook-shaped, whereas the head and nucleus of parasperm 2 was often less curved, more like that of eusperm (Figure 2G-I).

We conclude that *D. miranda* produces three sperm morphs – parasperm 1, parasperm 2 and eusperm. Parasperm 1 comprised 53% of the total sperm measured, parasperm 2 15%, and eusperm 32%.

#### 3.4.2. D. persimilis

HCA and GMM produced considerably differing results for *D. persimilis*. HCA identified two clusters – eusperm and parasperm. Sperm length of the single parasperm cluster identified by HCA was not normally distributed (W = 0.839, p < 0.001) indicating that the parasperm cluster does separate further into two subclusters, like that of *D. miranda* and *D. pseudoobscura*. GMM does identify these two parasperm subclusters but also identifies three eusperm clusters. These eusperm clusters overlap and are therefore unlikely to be distinct morphs, as suggested by normal distribution of HCA eusperm lengths (W = 0.991, p = 0.465).

Imaging of parasperm shows two distinct morphologies. Shorter parasperm 1 showed less coiling of the tail compared to medium parasperm 2, although parasperm 1 was often observed to have coiled back on itself, forming a loop. Both parasperm 1 and 2 were often observed to have a hook-shaped nucleus.

The two parasperm morphs were present in all individuals from which sperm were measured. We conclude that like *D. miranda*, *D. persimilis* produces three sperm morphs: short parasperm 1, medium parasperm 2, and long eusperm. Mean length of parasperm 1 was 45.3µm, parasperm 2 was 89.1µm, and eusperm was 299.1µm. Parasperm 1 comprised 20% of the total sperm measured, parasperm 2 11%, and eusperm 69%.

#### 3.4.3. D. pseudoobscura

*D. pseudoobscura* data was used to establish the parameters of the two models, as recently published data has shown strong evidence of three sperm morphs in this species (Alpern et al., 2019).

HCA identified three clusters, two parasperm and one eusperm. Eusperm length was normally distributed, after outliers were removed (W = 0.982, p = 0.094). Parasperm 1 length was normally distributed (W = 0.974, p = 0.185). Parasperm 2 length was normally distributed after outliers were removed (W = 0.990, p = 0.724).

GMM also identified these clusters but additionally separated eusperm into multiple clusters. As describe earlier for *D. persimilis*, these clusters overlap and are therefore unlikely to be distinct morphs. Both parasperm 1 and 2 clusters were present in all individuals measured.

Consistent with the descriptions by Alpern et al. (2019), we found that morphology is distinct between the short and medium clusters. Short parasperm 1 have a wider head and less coiling in the tail, compared to medium parasperm 2. We therefore conclude that, as has previously been described, *D. pseudoobscura* produce two parasperm morphs, and one eusperm morph. Mean length of parasperm 1 was 45.3µm, parasperm 2 was 89.1µm, and eusperm was 300.9µm. Parasperm 1 comprised 22% of the total sperm measured, parasperm 2 30%, and eusperm 48%.

### 3.5. subobscura – D. guanche and D. subobscura

#### 3.5.1. D. guanche

HCA and GMM gave different results for *D. guanche*. HCA identified two clusters, one parasperm and one eusperm. GMM identified four clusters, two parasperm and two eusperm. The two parasperm clusters are not distinct as those of the *pseudoobscura* species subgroup were and there was no clear distinction in morphology to support the designation of two parasperm morphs. Sperm length of the single parasperm cluster identified by HCA was normally distributed (W = 0.989, p = 0.358), further supporting a single parasperm cluster.

Of the eusperm clusters, the shorter appears, as with other species, to be outliers and broken eusperm. By HCA, eusperm length was not normally distributed (W = 0.923, p > 0.001), and was not normally distributed after removal of outliers (W = 0.977, p = 0.046). Visualisation of the data indicated eusperm length was marginally skewed, but not bimodal (S3).

We conclude that *D. guanche* produces two morphs. Parasperm mean length was 104.9µm, and eusperm mean length was 293.2µm. Parasperm comprised 48% of the total sperm measured.

#### 3.5.2. D. subobscura

Both HCA and GMM identified two clusters in *D. subobscura*. Both parasperm and eusperm lengths were normally distributed (parasperm W = 0.989, p = 0.508; eusperm W = 0.991, p = 0.821). Parasperm mean length was 103.7µm, eusperm mean length was 238.9µm. Parasperm comprised 57% *of the total sperm* measured.

## 4. Discussion

### 4.1. The *obscura* group species produce multiple sperm morphs

The D. obscura species group are the only sperm heteromorphic Drosophila species. All obscura group species previously studied have been found to produce a long sperm morph – the eusperm, and a short sperm morph – the parasperm. In this study, we re-measured total sperm lengths and nucleus lengths of eight obscura group species: D. affinis, D. azteca, D. bifasciata, D. miranda, D. persimilis, D. pseudoobscura, D. guanche and D. subobscura. We also found that all eight produce at least two distinct sperm morphs, and some species – D. miranda, D. persimilis and D. pseudoobscura – produce three.

We observed morphological differences between the eusperm and parasperm morphs. Parasperm were often more tightly coiled than eusperm and were often observed to have a hook-shaped nucleus. Some eusperm were also observed to have a bend in the nucleus, but this was less acute than that of parasperm. When imaged together, parasperm nuclei were often brighter than eusperm nuclei. As both morphs contain the same DNA content (Pasini et al., 1996), this indicates more condensed DNA in parasperm nuclei, as might be required in a smaller volume nucleus.

Future work could examine the differences in ultrastructure between eusperm and parasperm morphs, for example by electron microscopy (Pasini et al., 1996). This could also indicate developmental differences between morphs, such as in length, nuclear shaping and DNA condensation.

### 4.2. Sperm and nucleus lengths are consistent within species across studies

Mean sperm lengths for all species were consistent with previously published data. D. bifasciata eusperm mean length was slightly longer than previously published data (Joly et al., 1989), and D. guanche and D. miranda eusperm shorter than previously published data (Snook, 1997, Joly et al., 1989). However, all these species have been studied only once previously. For more commonly studied species, variability of mean sperm lengths is observed between studies and populations. It is likely therefore that this slight difference between data presented here and in previous studies reflects population and methodological variability.

We found no evidence of multiple parasperm morphs or ‘giant’ eusperm in D. subobscura, as was previously suggested in an early study by Beatty and Sidhu (1970).

### 4.3. D. miranda, D. persimilis and D. pseudoobscura produce three sperm morphs

D. pseudoobscura have recently been shown to have two parasperm morphs (Alpern et al., 2019). We wanted to assess whether its sister species, D. persimilis, and another closely related species, D. miranda, also produce two parasperm morphs, in addition to eusperm. We found that all three species of the pseudoobscura species subgroup produce three sperm morphs. The two parasperm morphs were distinct in length and morphology in all three species.

All other species were observed to have two sperm morphs, including the basal subobscura species subgroup, suggesting that this is the ancestral condition and the production of two parasperm morphs in the pseudoobscura subgroup is a derived trait. In future work, it would be of interest to investigate pseudoobscura subgroup-specific genes or gene duplications for their potential contribution to development of multiple parasperm morphs, for example by single cell RNA sequencing (Wei et al., 2024).

### 4.4. does *D. bifasciata* produce two eusperm morphs?

Cluster analysis raised the possibility of multiple eusperm clusters in D. bifasciata, although this clustering was inconsistent between hierarchical clustering and Gaussian mixture modelling. Multiple eusperm clusters may have been identified because of the large range of eusperm lengths observed in D. bifasciata. Other than length, no morphological differences were apparent. To establish whether eusperm of all lengths are capable of fertilisation, future work should measure sperm within fertilised eggs (Snook et al., 1994, Snook and Karr, 1998). Electron microscopy would indicate whether ultrastructural morphological differences are present between eusperm of differing lengths.

### 4.5. Function of non-fertilising sperm in sperm heteromorphic Drosophila

Although we have not investigated function in this work, it is likely that the morphs have the same or very similar function to those described in D. pseudoobscura: fertilising eusperm, and non-fertilising, protective parasperm (Holman and Snook, 2008). There is evidence that parasperm 2 also has a role in sperm competition in D. pseudoobscura (Alpern et al., 2019), and this may also be the case in D. persimilis and D. miranda, which also produce two parasperm morphs.

As all other obscura group species produce a single parasperm morph, it would be of interest to examine whether the single morph has both the protective and competitive functions, or just a single role. This would indicate whether the pseudoobscura subgroup, by evolving a second parasperm, has separated the two parasperm functions to specialised sperm morphs.

### 4.6. Cluster analysis as a method to identify sperm morphs

Previous studies have used kurtosis and ANOVA statistical methods to establish the number of sperm morphs produced by the obscura group species. Here we have used a different approach – cluster analysis – to predict the number of morphs produced, based on total sperm length and nucleus length data. We used hierarchical cluster analysis, with group number indicated by gap-statistic, and Gaussian mixture modelling. Gap-statistic guided hierarchical clustering was more conservative than Gaussian mixture modelling. For most species, the two models indicated different numbers of clusters. It was therefore necessary to also examine the clusters for morphological characteristics, in addition to the statistical approach.

## 5. Conclusions

Species of the obscura group produce multiple sperm morphs, a long eusperm morph, which is required for fertilisation, and one or two short parasperm morphs, which protect the eusperm in the female reproductive tract and may also function in sperm competition.

We have confirmed the presence of two parasperm morphs in D. pseudoobscura and have identified for the first time that its sister species, D. persimilis, and another closely related species, D. miranda, also produce two parasperm morphs.

## Data availability

All raw data is included as supplementary data.

## Ethics approval statement

Not applicable.

## Acknowledgements

We thank Drs. D. Bachtrog and T. Price for contributing D. miranda and D. pseudoobscura stocks, and the National Drosophila Species Stock Center (Cornell University, USA) and Kyorin-Fly Stock Center (Kyorin University, Japan) for other Drosophila species. We thank Drs. S. Christofides and W. Kay for their statistical analysis advice, and H. Bidwell, S. Patel, A. Roberts and J-L. Weston for their contribution collecting pilot data.

## Supplementary Data

**S1. Raw Data**

**S2. R Scripts for HCA and GMM Modelling**

**S3. Model Outputs**

**S4. Shapiro-Wilk Normality Test**

**S5. Parasperm length as a proportion of total sperm length**

**S6. Individual variation in sperm length – Sperm length by male scatterplots**

