## Supplementary data and code for "Clustering Sperm: A Statistical Approach to Identify Sperm Morph Numbers in the *Drosophila Obscura* Species Group": Supplementary3 Model outputs.docx

Supplementary 3: Model Outputs

### HCA

Hierarchical cluster analysis (HCA) outputs from hclust. **A** Optimal number of clusters by Gap Statistic modelling. **B** Dendrogram of clustering based on hclust HCA.

### GMM

Gaussian mixture modelling (GMM) output from mclust giving optimal number of clusters by GMM.

### *D. affinis*

#### HCA

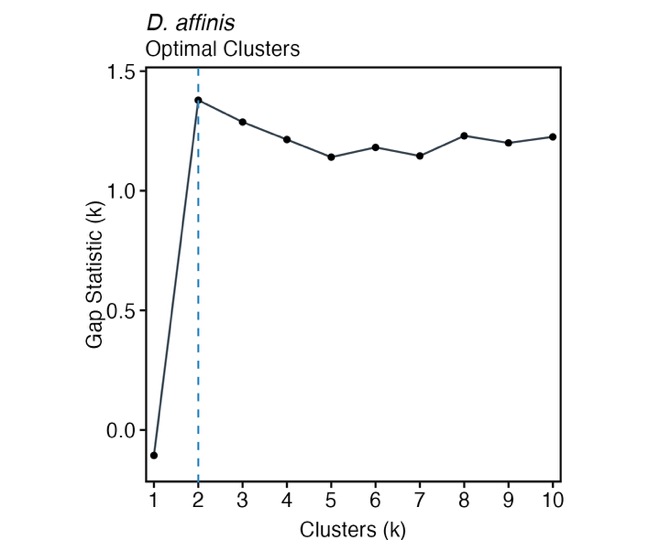

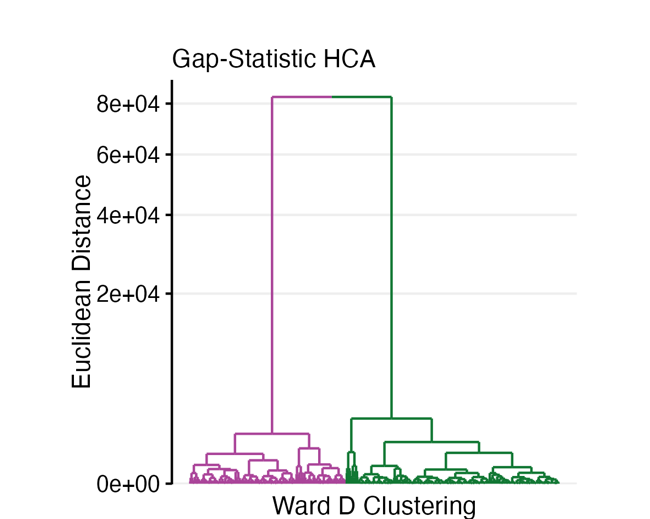

#### GMM

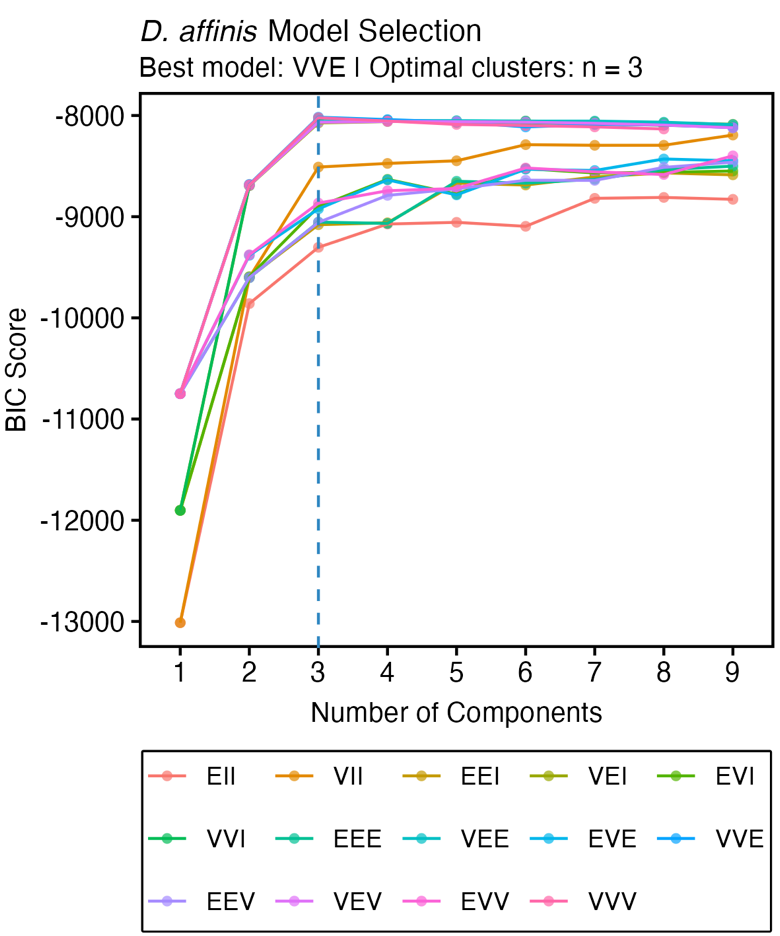

### *D. azteca*

#### HCA

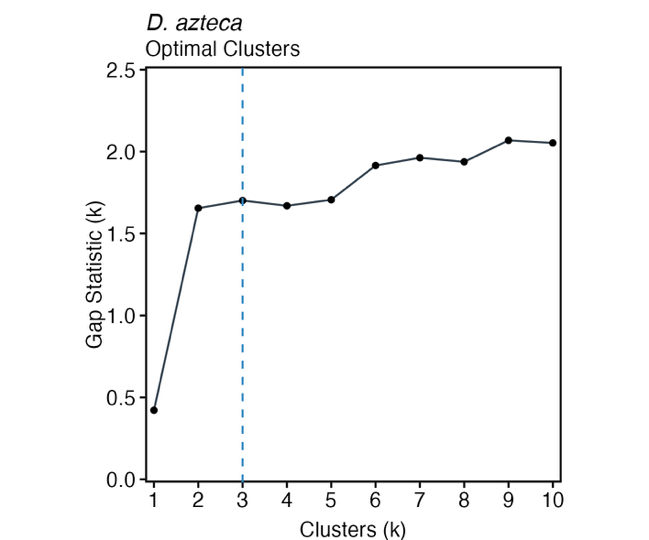

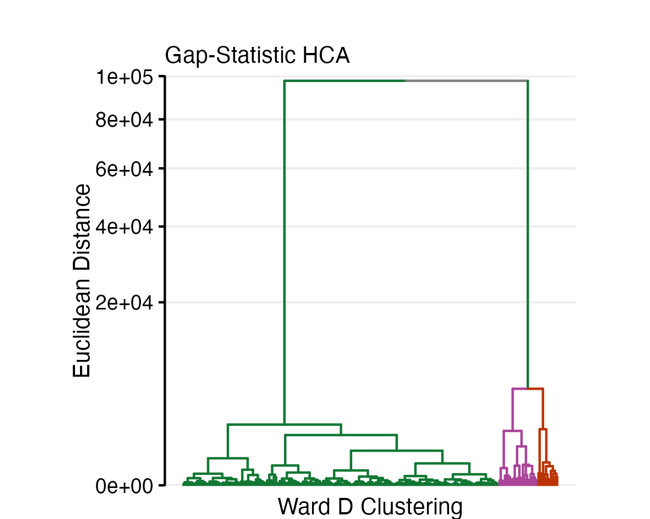

#### GMM

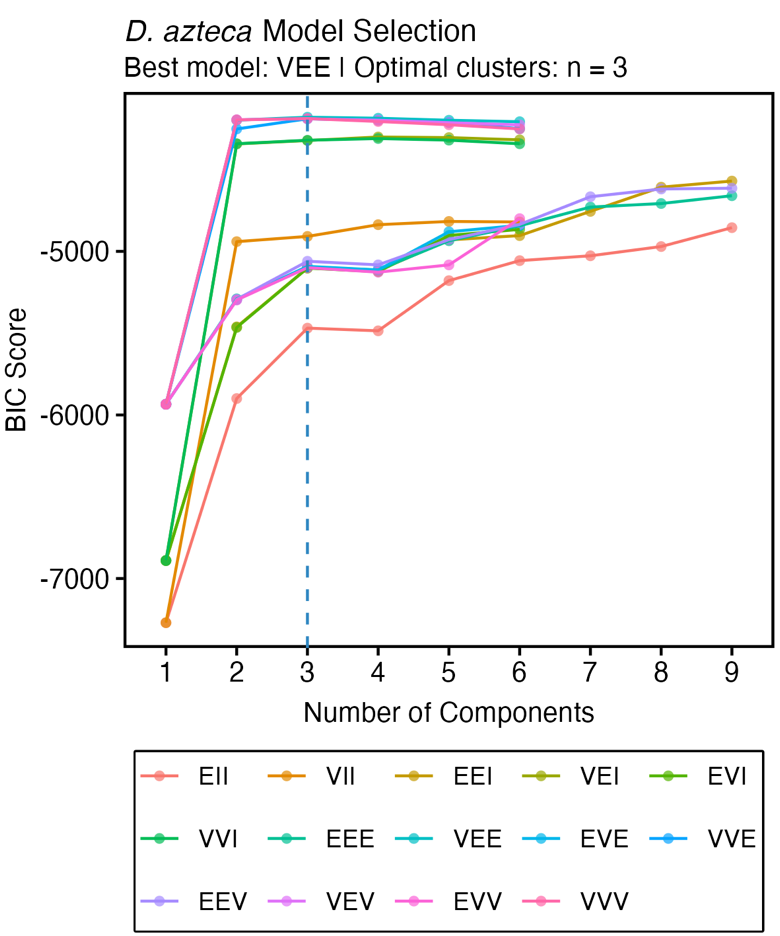

### *D. bifasciata*

#### HCA

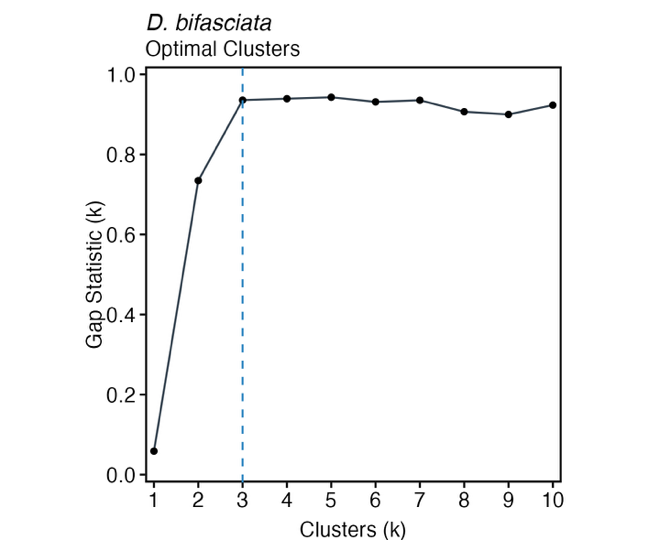

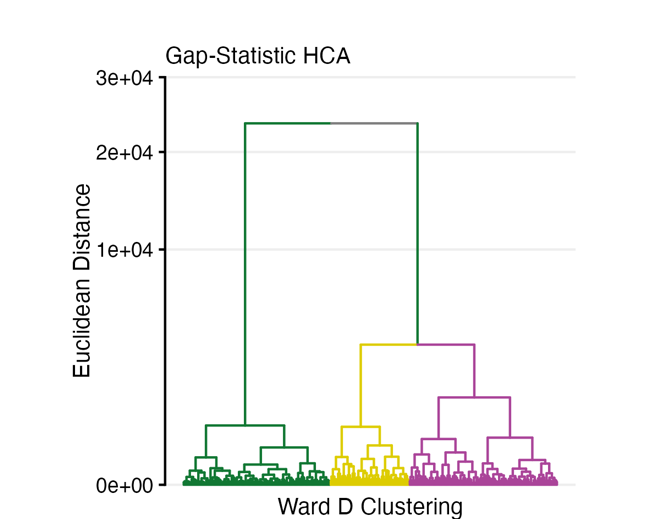

#### GMM

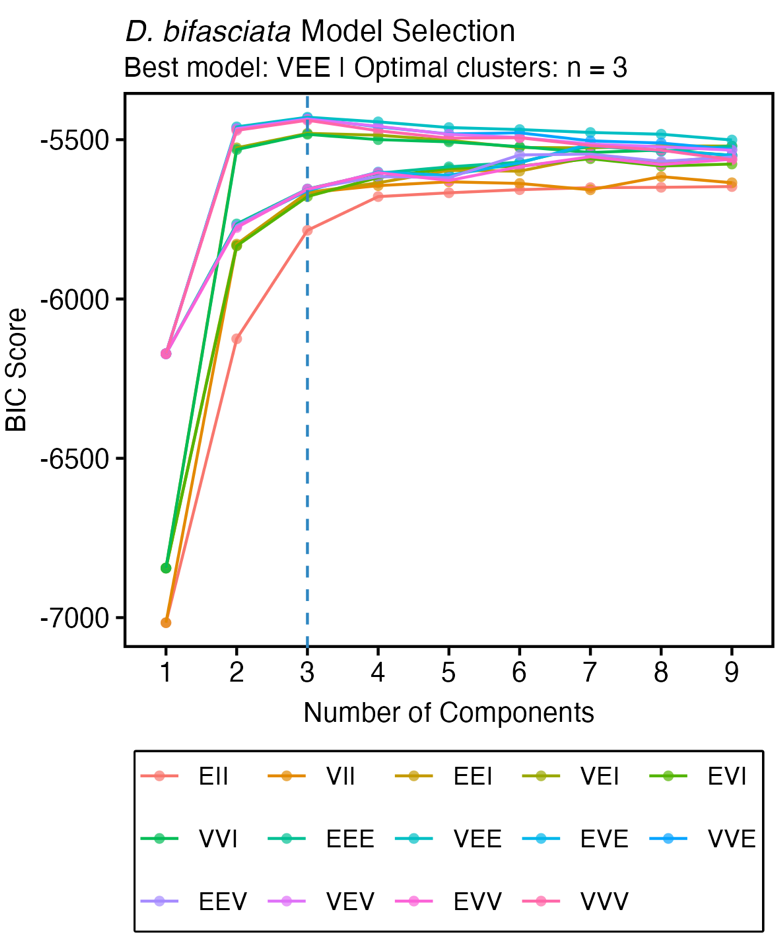

### *D. miranda*

#### HCA

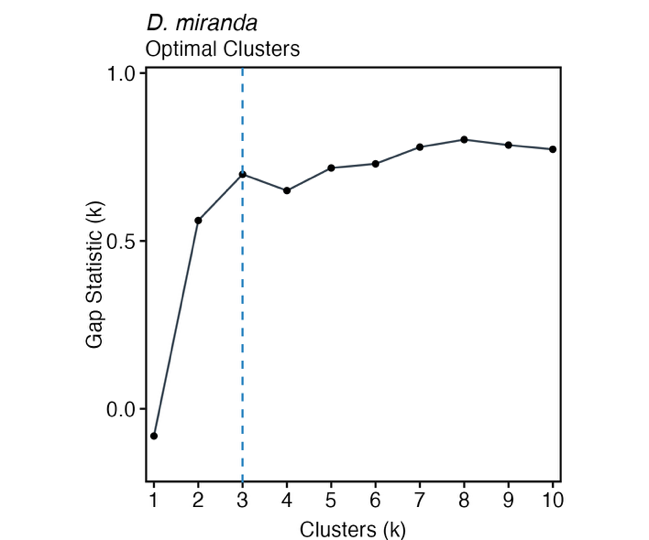

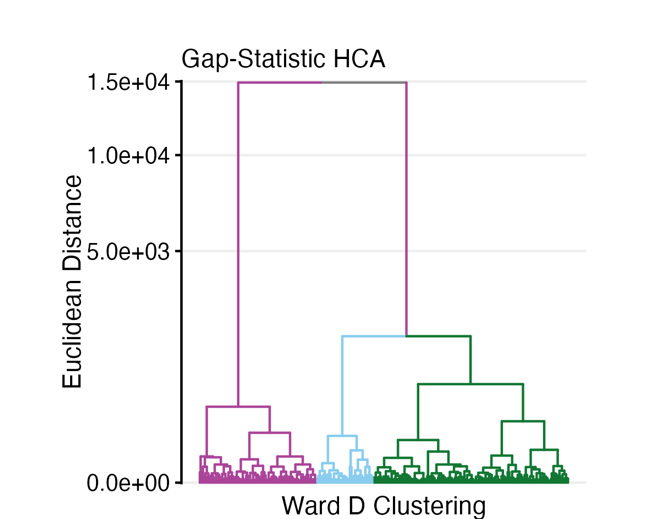

#### GMM

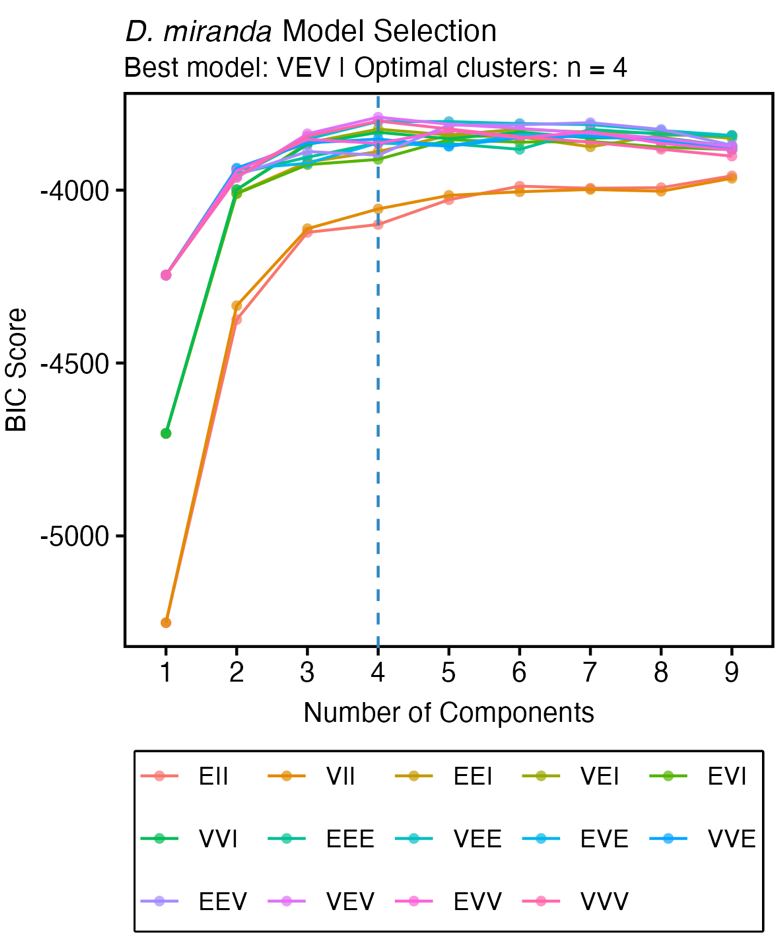

### *D. persimilis*

#### HCA

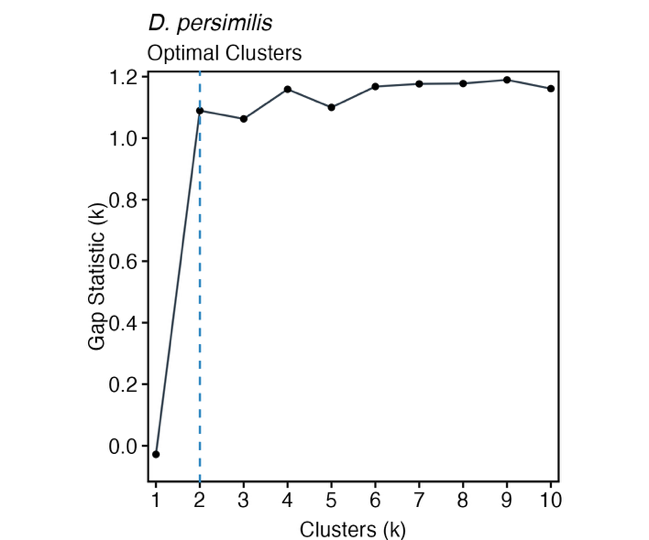

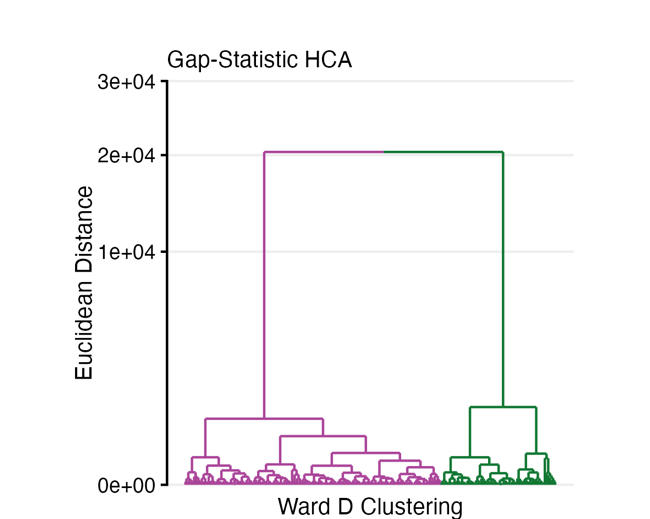

#### GMM

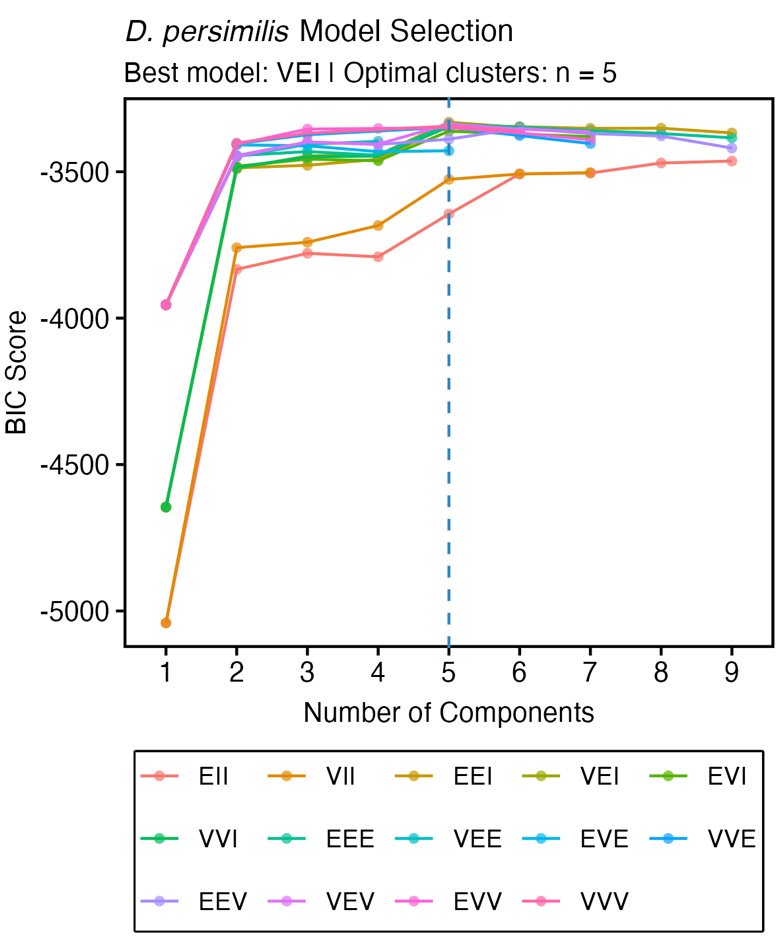

### *D. pseudoobscura*

#### HCA

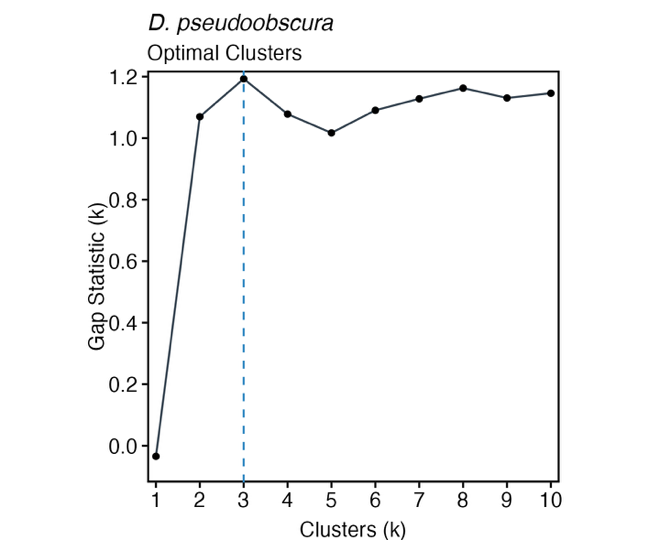

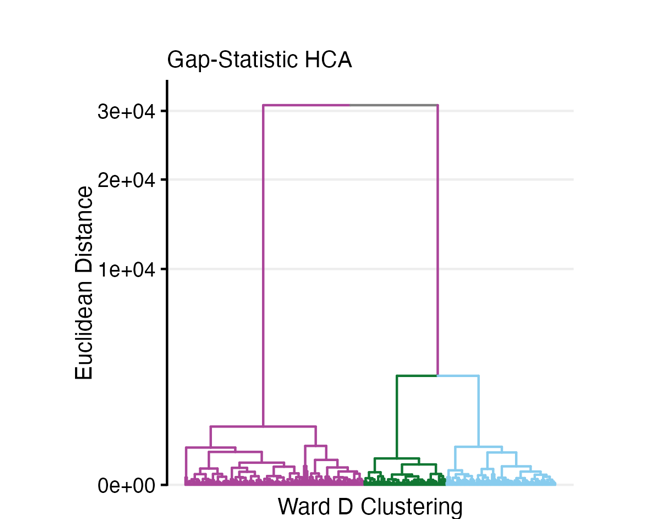

#### GMM

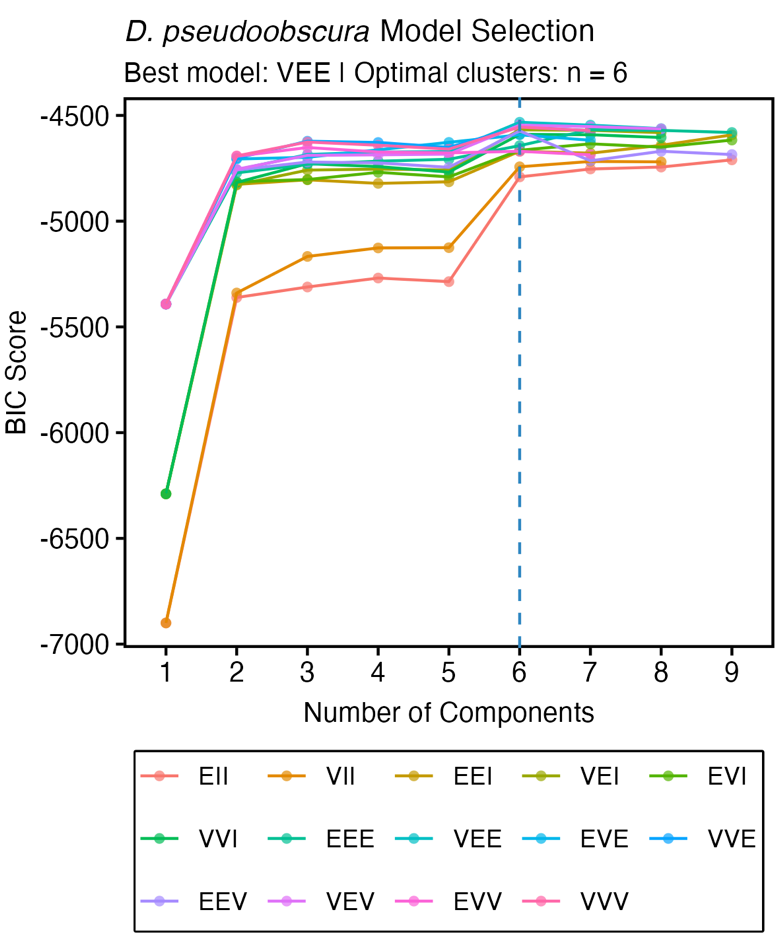

### *D. guanche*

#### HCA

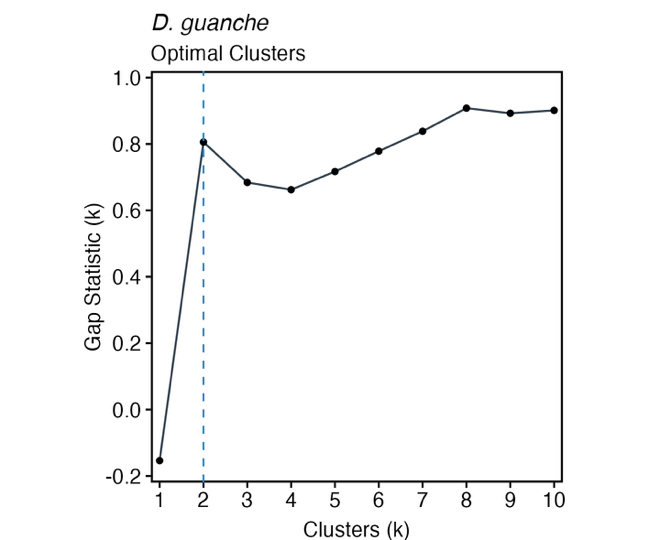

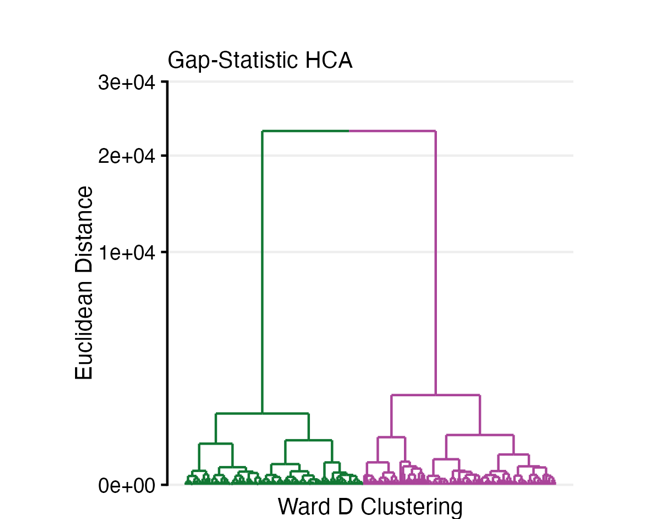

#### GMM

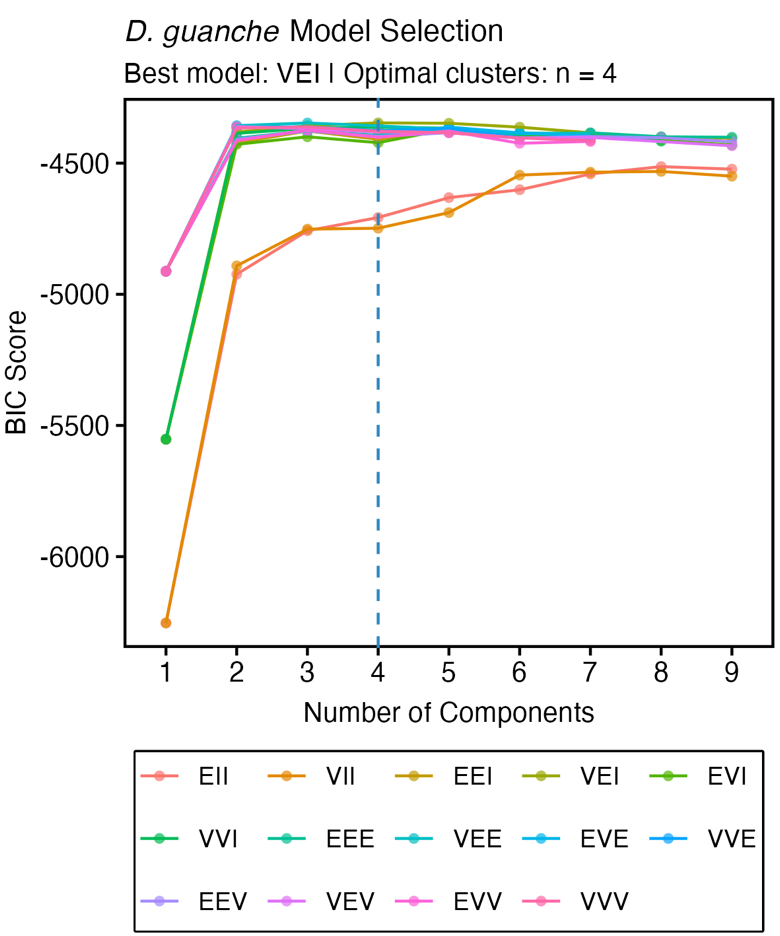

#### *D. guanche* eusperm density distribution

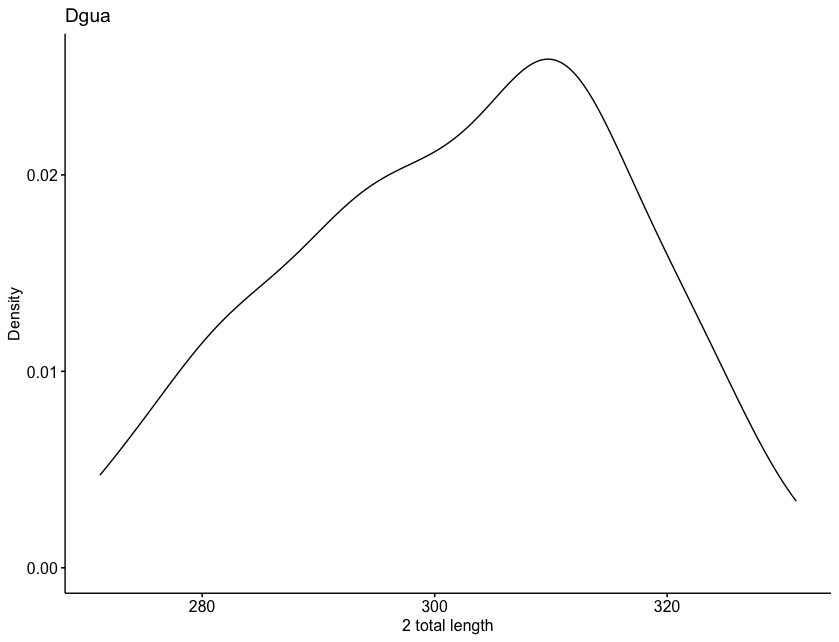

### *D. subobscura*

#### HCA

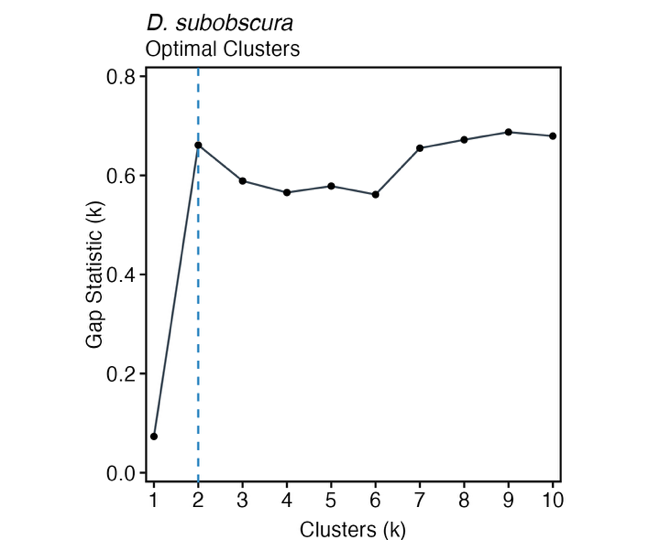

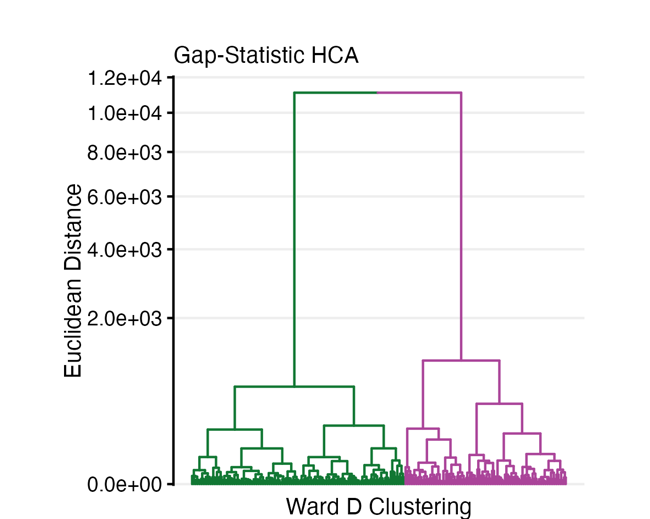

#### GMM

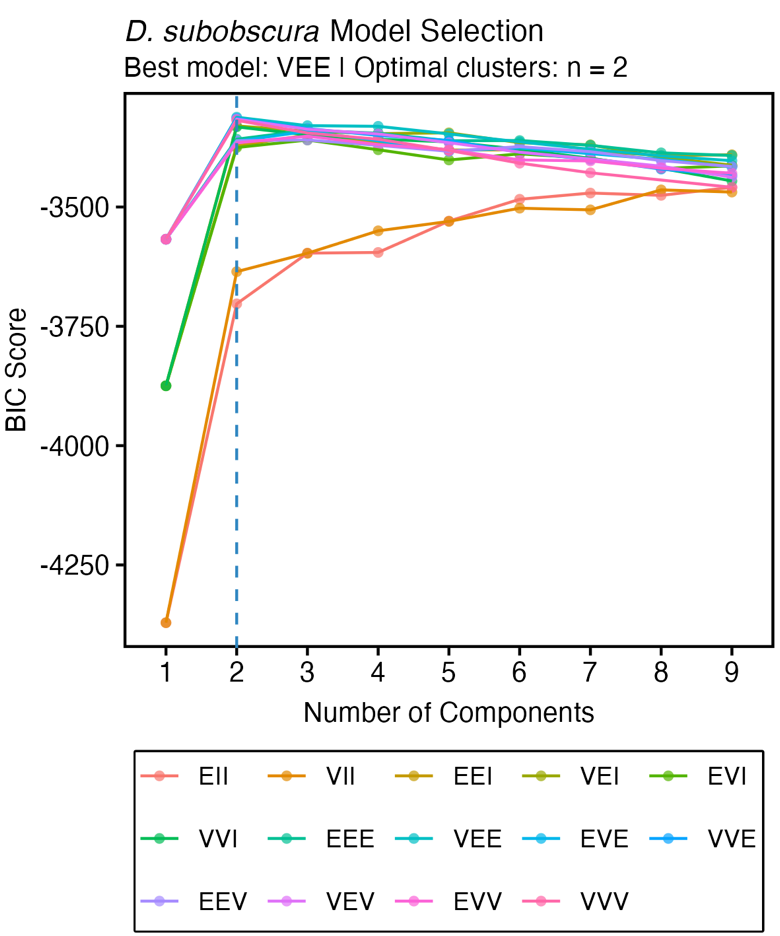
