## Supplementary figures and images for "Clustering Sperm: A Statistical Approach to Identify Sperm Morph Numbers in the *Drosophila Obscura* Species Group"

### Supplementary6 individual variation.docx

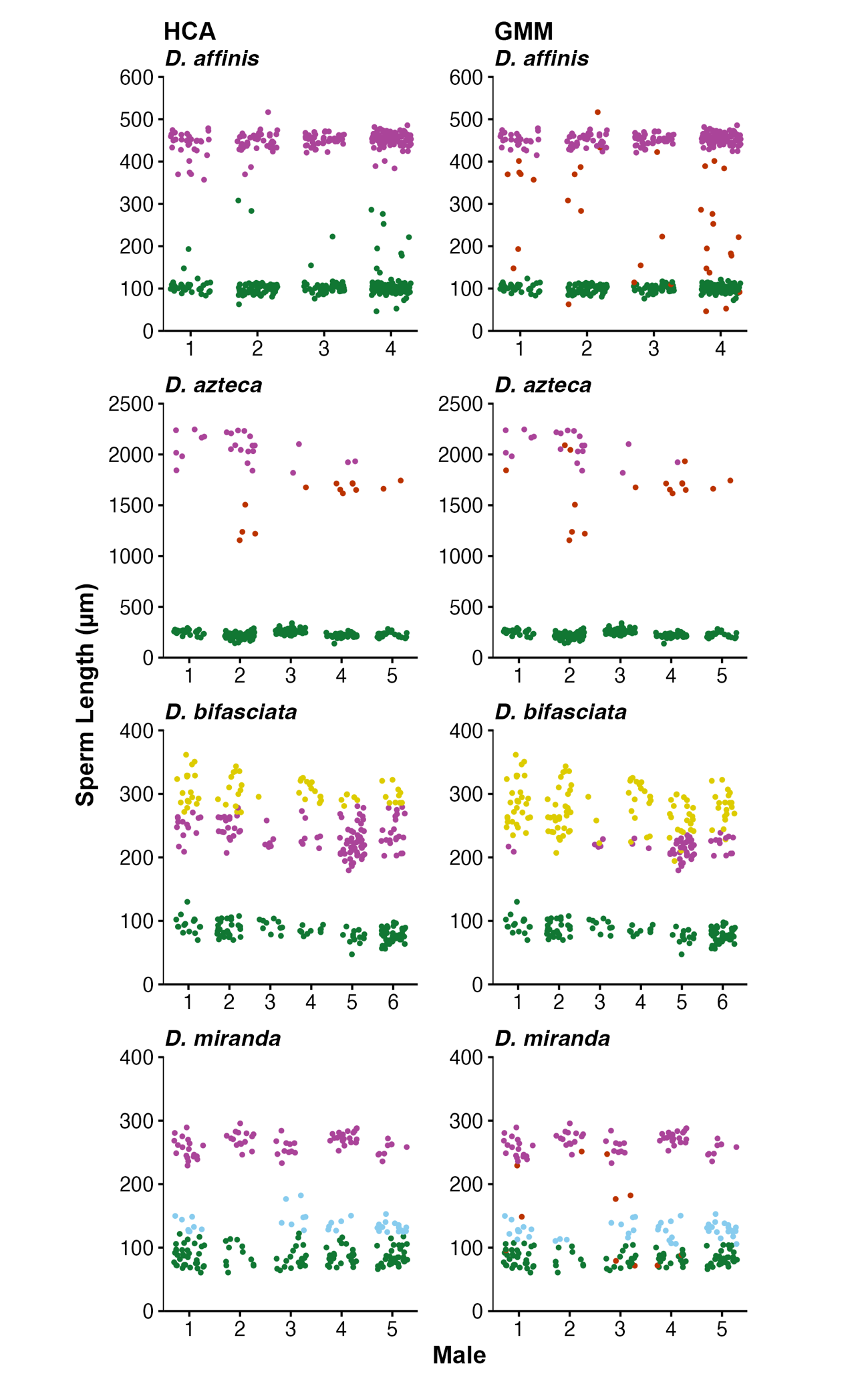

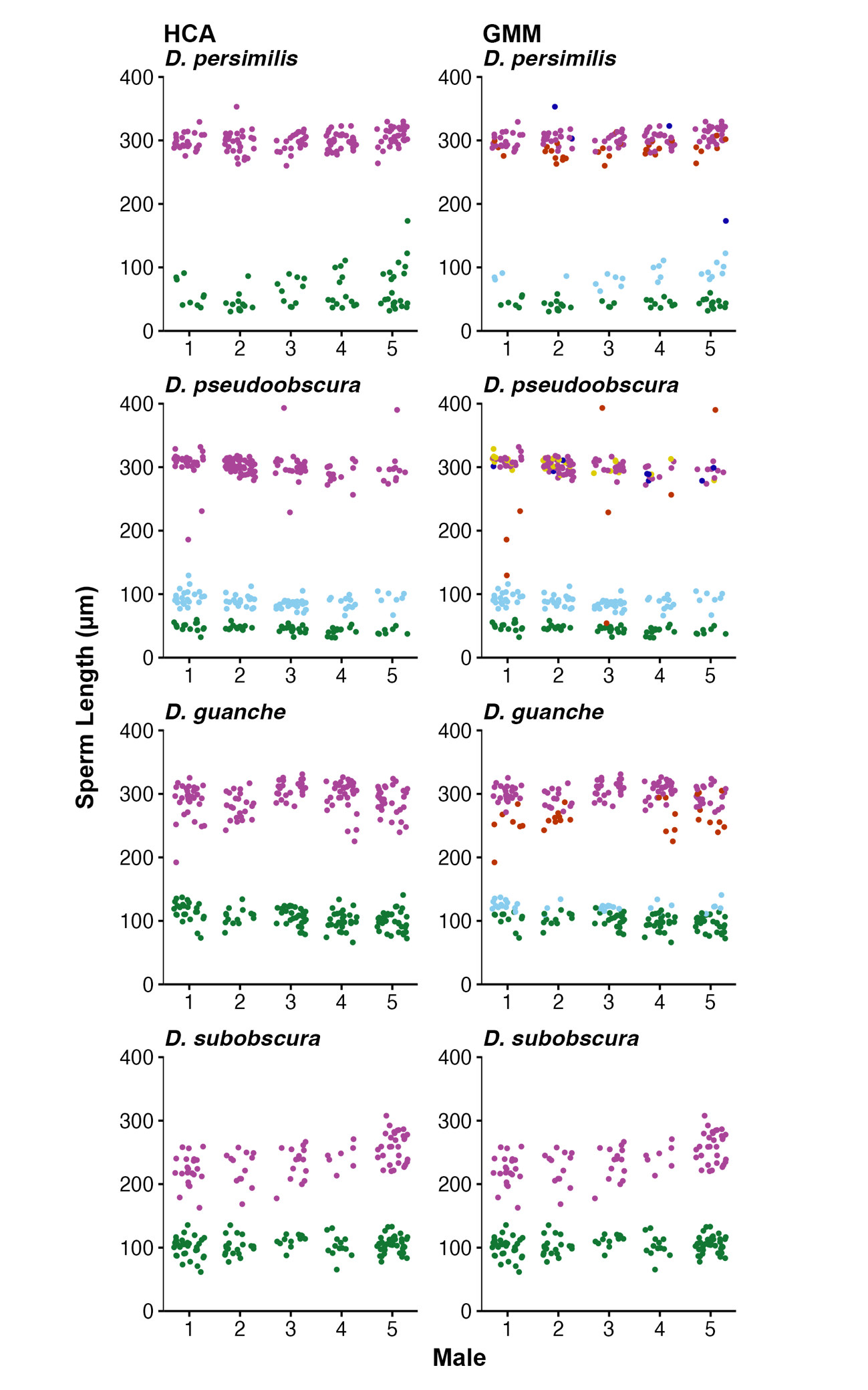
